# Polymer modelling unveils the roles of heterochromatin and nucleolar organizing regions in shaping 3D genome organization in Arabidopsis thaliana

**DOI:** 10.1101/2020.05.15.098392

**Authors:** Marco Di Stefano, Hans-Wilhelm Nützmann, Marc A. Marti-Renom, Daniel Jost

## Abstract

The 3D genome is characterized by a complex organization made of genomic and epigenomic layers with profound implications on gene regulation and cell function. However, the understanding of the fundamental mechanisms driving the crosstalk between nuclear architecture and (epi)genomic information is still lacking. The plant *Arabidopsis thaliana* is a powerful model organism to address these questions owing to its compact genome for which we have a rich collection of microscopy, Chromosome Conformation Capture (Hi-C), and ChIP-seq experiments. Using polymer modelling, we investigate the roles of nucleolus formation and epigenomics-driven interactions in shaping the 3D genome of *A. thaliana*. By validation of several predictions with published data, we demonstrate that self-attracting nucleolar organizing regions and repulsive constitutive heterochromatin are major mechanisms to regulate the organization of chromosomes. Simulations also suggest that interphase chromosomes maintain a partial structural memory of the V-shapes, typical of (sub)metacentric chromosomes in anaphase. Additionally, self-attraction between facultative heterochromatin regions facilitates the formation of Polycomb bodies hosting H3K27me3-enriched gene-clusters. Since nucleolus and heterochromatin are highly-conserved in eukaryotic cells, our findings pave the way for a comprehensive characterization of the generic principles that are likely to shape and regulate the 3D genome in many species.

## INTRODUCTION

In eukaryotic cells, the genome structure is characterized by a complex three-dimensional (3D) organization (1–3) that plays a crucial role in regulating gene function and expression (4, 5), and in determining cell-fate decisions (6–9), and cell-development (6, 10).

Microscopy and chromosome conformation capture (3C) (11) techniques have been used to unveil the architectural folding of the genome. Using microscopy techniques such as FISH (12), 3D-FISH (13, 14) and cryo-FISH (15), it was possible to visualize that each chromosome occupies a distinct portion of the nucleus called chromosome territory (CT) with a non-random radial location (12–14). High-throughput chromosome conformation capture (Hi-C) (16) confirmed the presence of CTs by probing much stronger cis- than trans-chromosome interactions (16, 17). Analysis of Hi-C maps also revealed the presence of a typical checkerboard pattern that reflects the physical segregation of the genome into multi megabase chromatin compartments (16, 18). These are characterized by different GC-content, gene density and epigenomic marks, suggesting that they mostly match the classical partition of the genome in hetero- and euchromatin (19–22). At the sub-megabase level, Hi-C experiments also revealed that the genome is organized in self-interacting regions termed topologically associating domains (TADs) (17, 23, 24), that have been also visualized by super-resolution microscopy approaches (25). In mammals and *Drosophila melanogaster*, TADs are considered the structural and functional units of the genome that define the regulatory landscape (26–28).

In animals, computational studies based on polymer modelling enabled to relate the organizational layers of the genome to specific active and passive physical mechanisms. The formation of chromosome territories may be facilitated by the slow relaxation dynamics of topologically-constrained polymers (29–32). The segregation between active and repressive chromatin compartments was suggested to be primed by micro-phase separation promoted by factors such as heterochromatin protein 1 (HP1) and polycomb group (PcG) proteins binding to repressive histone marks (33–37). The formation of TADs was associated with mechanisms of active (38, 39) or passive (40) loop-extrusion.

Within plants, *Arabidopsis thaliana* (*A. thaliana*) is an important model organism for structural genomics studies and has the most comprehensive collection of chromosome data available. Its genome, that is constituted by 5 chromosomes, is diploid, and is smaller (~120 mega-base pairs, Mbp) and gene-denser compared to other plant genomes, which makes it more suitable for computational studies.

In interphase, *A. thaliana* chromosomes are organized as well-defined chromosome territories (41–45) with generally no preferential radial positioning nor chromosome pairing (41, 46, 47). Notable exceptions are the short arms of acrocentric chromosomes 2 and 4, and the telomeres. The former host the nucleolus-organizing regions (NOR2 and NOR4), that typically associate in a single nucleolus at the centre of the nucleus (41, 48–50). The latter cluster at the nucleolar periphery (42–46). Within chromosome territories, *A. thaliana* chromatin contains active and repressive chromatin (51), that are organized in structural domains (42, 43). The chromocenters (including centromeres and peri-centromeres), that are the largest heterochromatic regions, are spatially and dynamically confined at the nuclear periphery (41, 46, 52, 53) and anchor protruding euchromatic loops of about 0.1-1.5 mega-base pairs (Mbp) resulting in a looped *rosette* overarching structure (46). In individual nuclei, chromocenters may also self-associate, leading to trans-chromosome contacts and larger heterochromatin foci (42–46). Hi-C data have also revealed the existence of long-range cis- and trans-chromosomal contacts, the so-called KNOT Engaged Elements (a.k.a. IHIs) structure (42, 43) that has structural counterparts in other plant species such as the compact silent centre (CSC) in rice (54). At fine scales, although TADs are not prominent features (44, 55), TAD-boundary-like elements have been shown to correlate with open chromatin and actively transcribed genes (44), and local self-interacting domains can be formed between H3K27me3-enriched gene-clusters (56, 57). At the scale of a few kilo-base pairs (kbp), local chromatin loops are suggested to connect the 5’ and 3’ ends of genes (45).

Previous modelling exercises (58), using a phenomenological ad-hoc coarse-grained polymer models, have suggested that the peripheral positioning of the chromocenters and the central localization of the nucleoli may be the consequences of entropic forces emerging from the formation of large permanent cis-chromosome loops and of steric constraints. However, a detailed description and characterization of the specific biological and physico-chemical mechanisms leading to the genome organization in *A. thaliana* is still missing.

Here, we test the hypothesis that the interactions between epigenomic states, as previously suggested (44, 45, 51, 59, 60) in other species, are driving forces of the genome structural organization in *A. thaliana* (61). We use polymer modelling and molecular dynamics together with epigenomic data to generate genome-wide chromosome models. By providing quantitative comparisons of our predictions with Hi-C and microscopy data, we demonstrate that four fundamental elements may be sufficient to determine the genome organization in *A. thaliana.* First, chromosomes need to be preconditioned as V-shaped objects. Second, the self-attraction of NORs shapes the overall nuclear organization. Third, the repulsion between constitutive heterochromatin and other epigenomic states explains the segregation and the localization at the nuclear periphery of chromocenters. Fourth, the self-attraction between facultative heterochromatin regions recovers the formation of self-interacting gene-clusters at local scales.

## MATERIAL AND METHODS

Our work consists of three main parts: (1) processing of ChIP-seq datasets to classify the *A. thaliana* chromosomes in epigenomic states; (2) genome-wide molecular dynamics simulations models, and (3) analysis of the obtained 3D models and validation against published experimental data.

### Epigenomic states analyses

ChIP-seq epigenomic data at 400 bp of resolution for 4 histone marks: H3K4me2, H3K4me3 (signatures of active genes), H3K27me3 (signature of facultative, polycomb-like heterochromatin), and H3K9me2 (specific to constitutive heterochromatin) were collected from published works (44, 45). To each 3kbp-genomic region of the genome, an average epigenomic signal was assigned for each of the 4 marks. A K-means algorithm (Euclidean distance, k=4) allowed clustering the 3kbp-regions into 4 groups: active chromatin (AC) enriched in H3K4me2/3, facultative heterochromatin (FH) enriched in H3K27me3, constitutive heterochromatin (CH) enriched in H3K9me2 and undetermined (UND) depleted in all these 4 marks. The lengths and the genomic localization of the domains assigned to each of the 4 chromatin states were used to generate the plots in **Figures 1B-C** and in **Supplementary Figure S1**.

**Figure 1.**
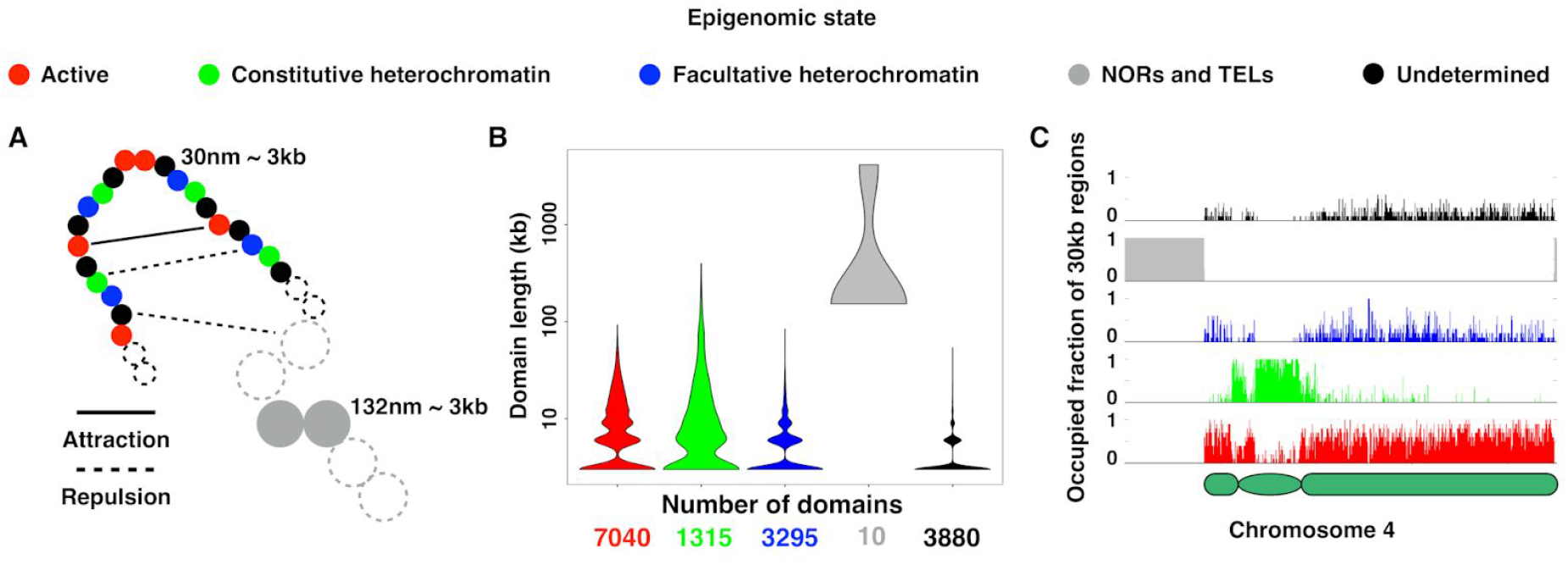
Epigenomics-driven co-polymer models of *A. thaliana* chromosomes. **(A)** Scheme of the polymer model. Chromosomes are modelled as self-avoiding bead-spring chains where each monomer represents a 3kbp-portion of chromatin and is characterized by its epigenomic state: active (red), constitutive heterochromatin (green), facultative heterochromatin (blue), NORs and telomeres (grey), and undetermined (black). Attractive or repulsive short-range interactions account for epigenomics-driven relationships between beads. **(B)** Distributions of the genomic length spanned by individual epigenomic domains (bottom: total number of domains per state). Each violin plot shows the density of the points smoothed with a Gaussian kernel with the standard parameters of R *geom_violin* function (133). **(C)** Profiles of each chromatin state along chromosome 4. Each profile is binned at 30 kbp and the height of the bars indicates the fraction of the 30kbp-bin occupied by 3kbp-regions of the corresponding chromatin state.

### Genome-wide chromosome simulations

#### The chromosome polymer model

Molecular dynamics simulations of the diploid *A. thaliana* genome were run using the 30nm-fibre model (62), in which each bead hosts 3 kbp of DNA sequence and has a diameter of 30 nm. Each *A. thaliana* chromosome was represented as a chain of beads using the Kremer-Grest bead-spring model (63, 64). This model allowed to represent chromatin as a 30 nm-thick fibre with a persistence length of 150 nm (65) and to avoid chain crossings (**Supplementary Methods**). The lengths of *A. thaliana* chromosomes (in bp and in models’ beads) and the genomic locations of special sequences (NORs and centromeres) were based on the reference genome TAIR10 (**Supplementary Table S1**).

Additional epigenomics-based short-range interactions were added to test how the attractions or repulsions between the *A. thaliana* chromatin states shape the chromosome organization. These interactions have been modelled using attractive or repulsive short-range (Lennard-Jones) potentials of variable strengths: from 10^−6^ to 1.0 k_B_T for repulsive interactions and from 0.025 to 1.00 k_B_T for attractive ones, where k_B_=1.38×10^−23^ J/K is the Boltzmann constant, and T=298 K is the temperature of the system (**Supplementary Methods**). Finally, the dynamics of the polymer model was simulated by integrating the (underdamped) Langevin equation of motion using LAMMPS (66). Each of the simulated trajectories lasted for 120,000 τ_*LJ*_ (internal LAMMPS time-unit) and allowed to generate 39 distinct models (1 model every 3,000 τ_*LJ*_ starting from 6,000 τ_*LJ*_). Since the equilibration time of dense or semi-dilute polymer solutions largely exceeded the simulation time (29), the obtained models are out-of-equilibrium structures in which physical properties of the system have reached a quasi steady state. See for example the **Supplementary Video S1**, that shows the convergence over time of the average contact probability (P) vs. the genomic distance (s).

#### Single-chromosome preliminary simulations

Single-chromosome simulations were performed for chromosome 4 (Chr4) described as a chain of 6,195 beads (excluding the NOR4 to simplify). To enhance the statistics, we simulated at the same time 5 copies of Chr4 placed in a cubic simulation box (size 2.76 μm) with periodic boundary conditions (DNA density of 0.004 bp/nm^3^ (29, 64)). In the initial conformation, each model chromosome was prepared in a linear rod-like shape to mimic an elongated mitotic state (29), and the 5 copies were placed in a random, yet non-overlapping arrangement. For each of the 50 simulated parameter sets (**Supplementary Figures S2-S4**), the dynamics of 10 independent trajectories were then simulated using LAMMPS (**Supplementary Methods** and **Figure 2**) and subsequently analysed (see below).

**Figure 2.**
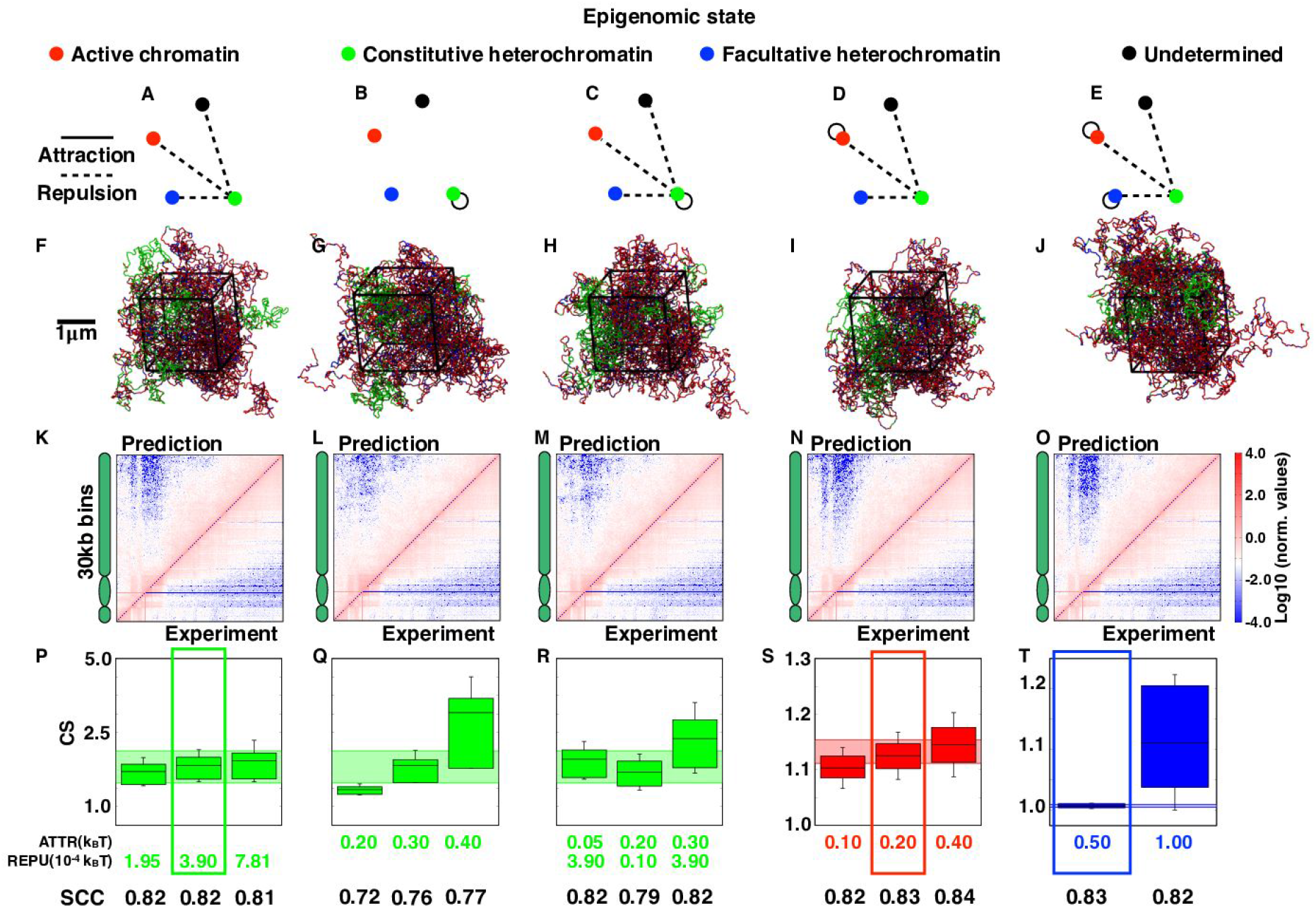
Parametrization of epigenomics-based interactions. **(A-E)** Networks representing the investigated sets of epigenomics-driven interactions. A link between two nodes represents a specific interaction. Other combinations are presented in **Supplementary Figures S2-S4**. **(F-J)** Illustrative snapshots of conformations obtained for each interaction network in the case of optimal consistency with the experiments (see panels **P-T**). Particles’ colours reflect their epigenomic state. These and other graphical representations of model chromosomes were rendered with the VMD graphical package (134). **(K-O)** Predicted contact maps for the six simulated interactions sets (top left triangles) are shown together with the cis-chromosome 4 Hi-C maps (bottom right triangles). **(P-T)** The compartment strength and the Spearman correlation analysis between the experimental and predicted contact maps (**Material and Methods**) are presented for a selection of parameters’ sets. Each Hi-C CS distribution is represented in the background of the plot with a colored band which spans the range of values from the first to the third quartiles. Boxes highlight the optimal cases.

#### The genome-wide models

For genome-wide simulations, we tested the effect of distinct initial conformations, which can be associated with mitotic-like chromosomes. Specifically, chromosomes were arranged as (i) linear rod-like objects as for the single-chromosome simulations (**Supplementary Figure S6A** and **Supplementary Video S2**), (ii) V-shaped objects generated by linear pullings along parallel directions (**Supplementary Figure S6B** and **Supplementary Video S3**), or (iii) V-shaped chromosomes generated by linear pulling along radial directions (**Supplementary Figure S6C** and **Supplementary Video S4**). V-shape cases mimicked the pulling of kinetochores by microtubules during metaphase and also accounted for the observation that the Hi-C interaction probability vs. the genomic distance (see **Figure 3E**) increases after 10Mb (the typical size of an arm in *A. thaliana* chromosomes) indicating an enrichment of inter-arm contacts. The procedures to generate the distinct chromosome shapes, to form the nucleolus by preconditioning the NORs arrangement, and to confine the chromosome models in a spherical nuclear environment are illustrated in the **Supplementary Videos S2, S3 and S4** and described in details in **Supplementary Methods**. For each of the tested mitotic-like chromosome shapes, 50 different initial conformations were simulated for the optimal parameter sets derived in the single-chromosome study (**Figure 3**). To test the relative importance of the model parameters, several variant genome-wide simulations were performed in which the optimal epigenomic-based interactions were perturbed one-by-one. The set of explored scenarios is summarised in **Supplementary Figures S7** and **S8**.

**Figure 3.**
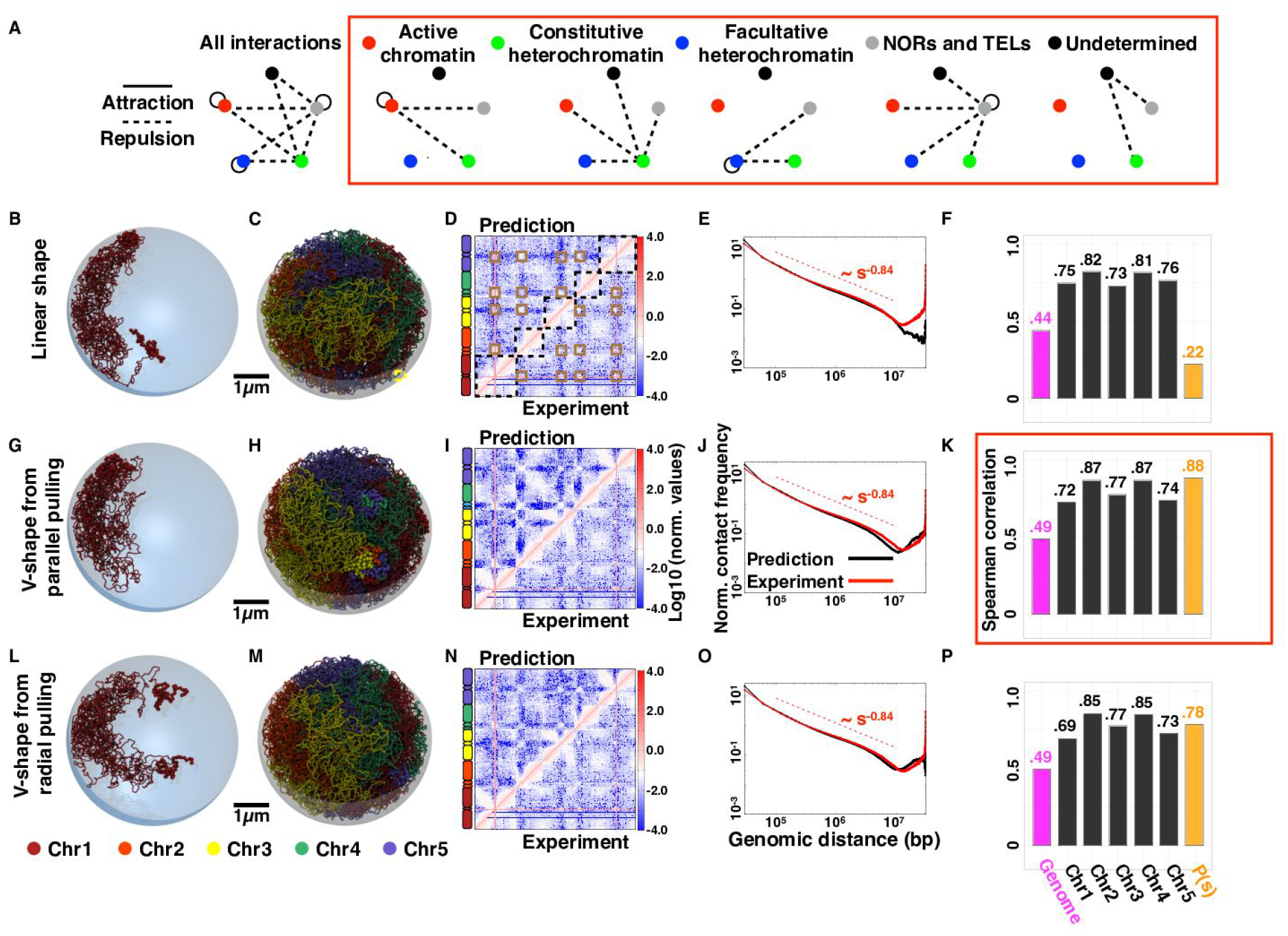
Genome-wide simulations: exploring the optimal epigenomics-based interactions and the effects of initial conformation. **(A)** Network of physical interactions used in the genome-wide simulations. In the red rectangle, the network is decomposed to highlight the interactions of each chromatin state. **(B-P)** Results of the genome-wide simulations for three alternative initial conformations: linear **(B-F)**, V-shape from parallel pulling **(G-K)**, and V-shape from radial pulling **(L-P)**. **(B**, **G**, **L)** Examples of initial (for a copy of chromosome 1) and **(C**, **H**, **M)** final (for all chromosomes) conformations. **(D**, **I**, **N)** Predicted genome-wide contact maps (top left triangles) together with the corresponding experimental Hi-C map (bottom right triangles). In panel D, the main features of the genome-wide contact maps are highlighted with overlying boxes: in black the cis-chromosome contacts accumulation, which is a signature of the presence of chromosome territories, and in brown the contacts between chromocenters. **(E**, **J**, **O)** Average cis-chromosome contact probability P vs genomic distance is computed from simulations (black curve) and from Hi-C (red curve). The power-low decay fitted on the Hi-C contact probability between 1 and 10 Mb is indicated with a dashed line. **(F**, **K**, **P)** Spearman correlation coefficients (SCC) computed to compare the models and the Hi-C data from genome-wide maps (magenta), cis-chromosome matrices (black), and P(s) (orange). The red box highlights the optimal case.

#### KNOT engaged elements-driven steered molecular dynamics

50 steered molecular dynamics simulations were performed to promote the spatial proximity between the KNOT engaged elements (KEEs) (42, 67) starting from the final snapshots of the optimal-model simulations. In particular, harmonics were applied between the central beads of the KEEs regions each spanning ~450 kbp. Briefly, the distances between the central beads of the KEEs were computed both *cis*- and *trans*-chromosome in all the 50 initial conformations and the closest bead pairs across all the snapshots were co-localized (**Supplementary Methods**) using the experimental FISH association rates (67) (20% of the KEE6-KEE1, 35% of the KEE5-KEE4, 66% of the KEE6-KEE3, and 16% of the KEE5-KEE10 pairs), or the average rate of 34% for all the other KEEs pairs.

### Analysis of the polymer models

#### Comparison with Hi-C data

Three Hi-C datasets for *A. thaliana* Col-0 seedlings were downloaded from the sequence read archive (SRA) (**Supplementary Table S2**) using *fastq-dump* (version 2.8.2, https://github.com/ncbi/sra-tools/wiki). Each experiment was processed through the TADbit pipeline (68) (https://github.com/3DGenomes/tadbit). Briefly, the pipeline consists of (i) Checking the quality of the FASTQ files; (ii) Mapping of the paired-end reads to the A. thaliana reference genome (release TAIR10 ftp://ftp.ensemblgenomes.org/pub/release-40/plants/fasta/arabidopsis_thaliana/dna/) using GEM (69) taking into account the DpnII restriction enzyme cut-site using fragment-based mapping (68); (iii) Filtering to remove non-informative reads using the following (default) TADbit filters: self-circle, dangling-ends, error, extra-dangling-ends, duplicated and random-breaks (68); (iv) Merging of the datasets into a single one for the restriction enzyme DpnII; (v) Normalization of the merged datasets using the OneD (70) method at 3 kbp and 30 kbp resolution. Before merging the datasets (point (iv)), their mutual consistency was verified using the reproducibility score (R-score) (71). The obtained R-score values ranged between 0.56 and 0.84 indicating consistency between the merged datasets (71).

For each simulated configuration, contacts were computed using a cut-off distance of 200 nm which characterizes roughly the spatial resolution of Hi-C experiments (72, 73). From these contacts, we then built contact maps for the single-chromosome and genome-wide cases. From these maps, the average contact probability P(s) was quantified by averaging the predicted contact frequency over all the pairs of loci at the same genomic distance (s). To compare visually predicted and experimental Hi-C data, the number of contacts in the models were re-scaled such that the average number of contacts at a genomic distance of 300 kb (the number of bp in one Kuhn segment of the polymer models) equals the experimental one. Quantitative comparisons were made using the Spearman correlation coefficient (SCC) analysis applied to the P(s), genome-wide and cis-chromosome matrices, and the compartment strength (CS) analysis (74) for each chromatin state. CS definition is based on the observed-over-expected (OoE) map. This map is computed as the entries of the contact or Hi-C map divided by the P(s) for the corresponding genomic distance s. For each bin (b) of a given epigenomic state, the CS was then quantified as the ratio between the average OoE value of b with bins of the same epigenomic state and the average OoE value of b with any other bin in the same chromosome. CS scores of all the bins of a given epigenomic state are then pulled together to form the CS distribution of that state, that were shown in the figures of this work. In this metric, no compartmentalization corresponds to CS=1, whereas any pattern of compartmentalization yields to CS>1.

#### Nuclear positioning of genomic regions

Radial positions of the beads for each epigenomic state were used to build the histogram of the number of beads per concentric nuclear shell of width 250 nm. Similar conclusions can be obtained using binnings of 125 or 500 nm (**Supplementary Figure S9**). The probabilities per shell were obtained by dividing the number of particles by the volume of each shell. For each parameter set, histograms were computed for each of the replicates. The means and standard errors computed over the replicates are reported as bar plots and error bars respectively (**Figures 4F-J**; **Figures 5G-L**; **Figures 5G-I**; **Supplementary Figure S9**; and **Supplementary Figures S10-S17 panels A-E**). To compare the radial distributions of two parameter sets, Wilcoxon tests were performed to compare each paired (corresponding replicates) distribution without assuming any specific shape for the two distributions under comparison. The null hypothesis is that the mean heights over the replicates give zero difference and the alternative is that the difference of the means is either higher or lower than zero (two-sided statistical test). A very stringent threshold for significance (p-value<0.0001) was chosen to single out only the most relevant differences in bins occupancies.

**Figure 4.**
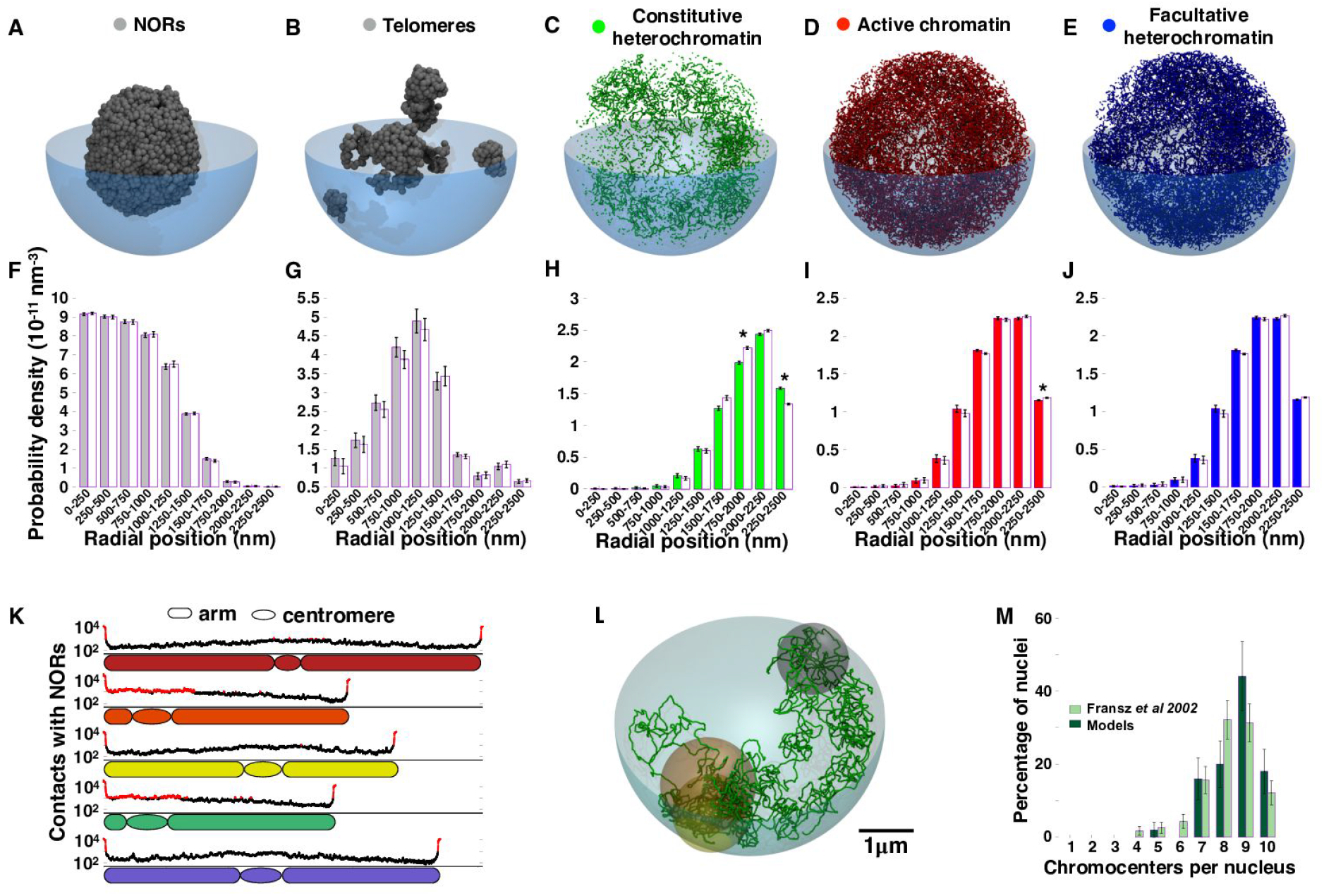
Nuclear positioning of the epigenomic states. **(A-E)** Representative snapshots of the nuclear localization of 3kbp-beads for each epigenomic state. **(F-J)** Probability to find a bead in concentric shells (thickness 250 nm) of the nucleus for each epigenomic state for the optimal interaction model (**Figure 3A**) (coloured bars). White bars illustrate the results of a null model (**Supplementary Figure S7B**) where all but NORs and telomeres interactions are switched off. The significantly enriched/depleted shells with respect to the null model (Two-sided Wilcoxon statistical test with p-value<0.0001) are marked with asterisks. Similar conclusions can be derived by varying the thickness of the shells to 125 or 500 nm (**Supplementary Figure S9**). **(K)** Predicted number of contacts within 200 nm with NORs particles along the different chromosomes. The top 10% of the contacting regions are highlighted in red. **(L)** Each group of centromeric beads per chromosome is represented by a sphere (radius = the radius of gyration of the constitutive particles; centre = their centre of mass). Spheres with a volume overlap larger than 34% the volume of the smaller sphere are part of the same focus. **(M)** Number of distinct centromeric foci per simulated conformation (dark green bars) and per experimental single cell (46) (light green bars). Error bars were computed as the square root of the average value under the hypothesis of a Poissonian distribution. For each bin, we tested if the predicted average frequency is similar to the observed experimental counts (null hypothesis) by computing the p-value of the predictions assuming Poisson distribution for experiments. All p-values were higher than 0.02 making impossible to reject the null hypothesis.

**Figure 5.**
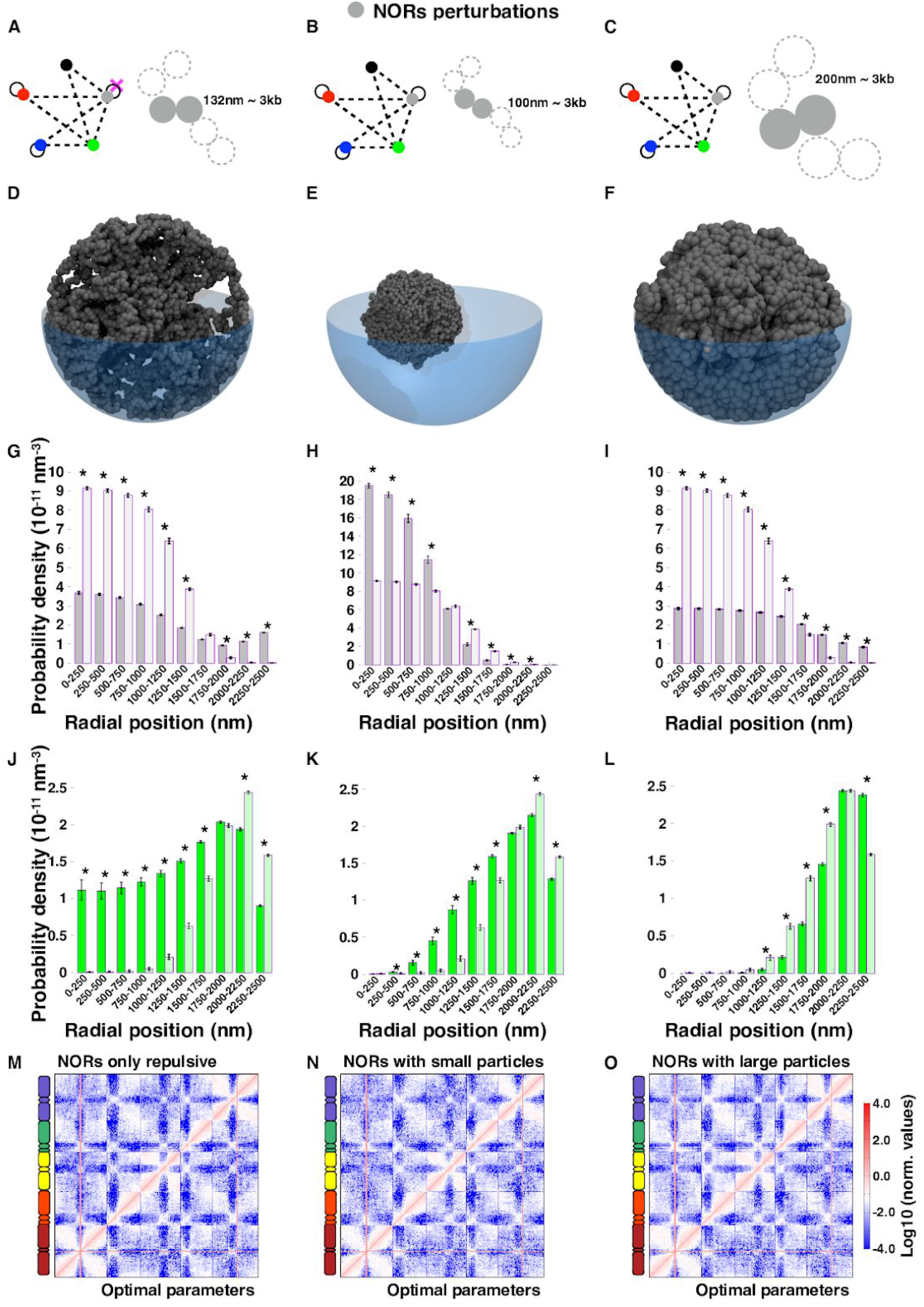
NORs shape the *A. thaliana* nuclear organization. **(A-C)** Perturbed interaction networks. The interactions that are removed from the optimal case are crossed in magenta. The cartoons of the polymers illustrate the variation in size of the NORs beads. **(D-F)** Illustrative snapshots for NOR beads in each of the perturbed systems shown in panels A-C. **(G-L)** Distributions of the radial positions of the 3kbp-regions in the perturbed systems (dark colour) compared to the optimal interaction model (light colour) for the NORs (G-I) and the CH (J-L) (epi)genomic states. Significant differences (two-sided Wilcoxon test p-value<0.0001) are marked with asterisks. **(M-O)** Genome-wide contact maps for the perturbed cases (top left triangles) vs maps obtained from the optimal interaction model (bottom right triangles). Complementary results on the simulations presented here are shown in **Supplementary Figures S12-S14**.

#### Number of distinct chromocentric regions

Each chromocenter was considered as a sphere (radius = the radius of gyration of the constitutive particles; centre = their centre of mass) (**Figure 4L**). The overlap between two spheres was computed using the formula in Weisstein, Eric W. “Sphere-Sphere Intersection.” From MathWorld--A Wolfram Web Resource. http://mathworld.wolfram.com/Sphere-SphereIntersection.html. The average number of distinct chromocenters was computed varying the overlap volume threshold between 0.0 and 1.0 every 0.01. The threshold of 0.34 corresponded to an average of 8.6 distinct chromocenters, which was the closest to the experimental measure of 8.6+/−0.2 chromocenters (46).

## RESULTS

### Epigenomics-driven folding of *A. thaliana* chromosomes using polymer models

In this study, we tested the hypothesis that specific interactions mediated by epigenomics are driving forces of the genome structural organization in *A. thaliana*. To this end, we applied to a plant species a modeling method that has been previously applied to the fly (33, 37) and human (35, 36, 75–79) genomes and that posits that epigenomics-driven interactions partition the genomes into spatial compartments. Here, we built polymer models of the chromosomes using beads of 30 nanometers (nm) hosting each 3 kilobase pairs (kbp) of chromatin (**Figure 1A**). We used short-range attractive and repulsive interactions between 3kbp-regions defined by their epigenomic state in conditions of density and confinement mimicking the nuclear environment.

Based on the enrichment in epigenomic marks (44, 45), we assigned to each 3kbp-region one of four chromatin states (33) (**Material and Methods**). A total of 52% of the genome was assigned to active chromatin (AC) enriched in H3K4me2 and H3K4me3, 14% to facultative, polycomb-like heterochromatin (FH) decorated with H3K27me3, 14% to constitutive heterochromatin (CH) enriched in H3K9me2, and 12% to undetermined (UND) regions that were depleted in all the considered epigenomic marks (**Figures 1B** and **C**, and **Supplementary Figure S1**).

Extending previous modeling approaches, the remaining parts of the *A. thaliana* genome were assigned to two other genomic categories that we used as complementary to the epigenomic states. The nucleolar organizing regions (NORs), that are the constitutive sequences of the nucleolus (41, 49, 50), account for 6% of the genome and are localized in the small arms of chromosomes 2 (NOR2) and 4 (NOR4). The telomeres, that account for about 1% of the genome, are the ~150 kbp regions at the ends of each chromosome (**Figure 1C**, **Supplementary Figure S1**, and **Supplementary Table S1**). To account for the high concentrations of RNA and proteins within the nucleolus (41, 49, 50) and for the typically increased stiffness of the telomeres (80), NORs and telomeric 3kbp-regions were modelled as thicker beads (**Figure 1A)** of diameter 132nm allowing to mimic the formation of a spherical nucleolus of radius about 1.4 μm as typically observed in *A. thalian*a nuclei (50, 81) (**Supplementary Methods**). Excluding the NORs, the more extended epigenomic domains of ~400 kbp were assigned to constitutive heterochromatin (CH) (**Figure 1B**) and, consistently with previous analysis (59, 60), were localized at the chromocenter (centromeric and pericentromeric region) of each chromosome (**Figure 1C** and **Supplementary Figures S1A-E**). In our analysis, chromocenters are interspersed with short sequences hosting active chromatin (maximum size 54 kb) and facultative heterochromatin (maximum size 18 kb), which is consistent with the presence of ‘punctual’ active genes like the 5S rRNA gene clusters in the peri-centromeres of chromosomes 4 and 5 (82). Overall, the genomic regions occupied by active chromatin and facultative heterochromatin were organized into many interspersed domains of shorter lengths (maximum 93 kbp for AC and 84 kbp for FH, and median 6 kbp for AC and 3 kbp for FH).

### Single-chromosome analysis allows parametrizing the epigenomics-driven interactions

To find the optimal epigenomics-driven model, we designed a strategy to minimize the amount of calculations needed and yet get a good understanding of how the different parameters affect the final compliance with the experiments. We focussed initially on the folding of chromosome 4 as it is the shortest in *A. thaliana* (**Supplementary Table S1**) and neglected for simplicity the nucleolar organizing region (NOR4). We simulated a toy-model with 5 copies of chromosome 4 with random initial placements and orientations in a cubic box with periodic boundary conditions at the typical DNA density (ρ=0.004 bp/nm^3^) of the *A. thaliana* nuclear environment (29, 64). In the initial conformation, the 5 model chromosomes were prepared as linear rod-like objects to mimic elongated mitotic states (29) (**Material and Methods**).

As illustrated in **Figures 2A-E**, we tested different types of short-range interactions involving the epigenomic chromatin regions defined above: self-attraction (full circles) and repulsion (dashed lines). Unless specified, the model beads interact with an excluded volume potential that allows the fibre to maintain a thickness of 30 nm and to avoid chain crossing (**Supplementary Methods**). Specifically, we started exploring three distinct scenarios for the interactions involving the constitutive heterochromatin (CH) regions (**Figure 2**): repulsions between CH beads and the other epigenomic states (**Figure 2A**), self-attraction between CH beads (**Figure 2B**), and a combination of the two (**Figure 2C**). These cases aimed to describe interactions driven by heterochromatin protein 1 (HP1) both *in vitro* and *in vivo*. Recent papers showed that HP1 (or its counterparts in plants LHP1 and ADCP1) can promote liquid-liquid phase separation (LLPS) mediated by intrinsically disordered regions (IDR) *in vitro* (83, 84), can favour the formation of heterochromatin compartments *in vivo* (83, 85, 86), and generate a phase from which some proteins may be excluded (83, 84). Overall, the collected evidence is yet not conclusive on whether the *in vivo* formation of HP1 phases is mainly driven by attraction between HP1 or by repulsion of HP1 with other chromatin states or with other chromatin-binding proteins. LLPS, in general terms, can be triggered via both self-attraction of a compound (in this case HP1) or via a repulsive interaction of a compound with the others present in the system, as it occurs for hydrophobic phases (87). On top of the CH-driven interactions, we tested separately the self-attraction of the active-chromatin (AC) regions (**Figure 2D** and **Supplementary Figures S2-S4**) and of the facultative heterochromatin (FH) beads (**Figure 2E** and **Supplementary Figures S2-S4**). The latter interactions are based on the observations that the binding of RNA PolII to active genes can prime micro-phase separation (88, 89) and polycomb-mediated interactions can form local domains hosting H3K27me3-enriched gene-clusters (56, 90, 91).

In total, we tested 50 distinct scenarios with strengths of interaction ranging from 10^−6^ to 1.00 k_B_T for repulsions and 0.025 to 1.00 k_B_T for attractions (**Supplementary Figures S2-S4**). One k_B_T typically represents an energy of the order of the thermal noise. The selected values of interaction strength allowed sampling distinct scenarios in which each of the imposed interactions varied between a small fraction to the same order (1.00 k_B_T) of the thermal noise. For each of the 50 distinct scenarios, we simulated 10 independent trajectories of a few hours (30, 64) using Langevin dynamics (**Material and Methods**) and obtained an ensemble of ~400 conformations per parameter set (**Figures 2F-J** for illustrative snapshots at the end of the simulations).

To select the optimal interaction model, we computed predicted contact maps at 30 kbp resolution (**Figures 2K-O** and **Material and Methods**) on the generated 3D models and compared them with the experimental Hi-C maps (44, 45). The comparisons included the Spearman correlation coefficient (SCC) (92, 93), which captures the overall similarity between model and experiment, and the compartment strength (CS) (74) for each epigenomic state (CH, AC, and FH), that quantifies the degree of compartmentalization of a given state by measuring the contact enrichment within versus between chromatin domains (**Figures 2P-T**, **Supplementary Figures S2-S4**, and **Material and Methods**).

The obtained SCC values were larger than 0.70 for all the tested interaction models (**Figures 2P-T** and **Supplementary Figures S2-S4**) indicating that the explored models allow capturing the overall arrangement of chromosome 4. These high correlations are mainly due to the formation of a segregated domain (**Figures 2K-O**) at the chromocenter, that, in the models, is favoured by the CH-driven interactions. In fact, the addition of self-attraction within AC or FH improves marginally the SCC only by 1%. The difference between CH-scenarios (**Figures 2P-R**) is also weak (at most 0.05) with the CH repulsion scenario overall leading to higher similarity between models and experiments.

As expected, as the strengths of epigenomic interaction are increased, the CS augments (**Figure 2P-T** and **Supplementary Figures S2-S4**). Interestingly, we found that optimal parameters for CS are also close to optimal for SCC, with weak or intermediate values for interaction strengths. For example, in the simulations where the optimal CH self-attraction (E_CH_=0.30 k_B_T) is combined with increasing AC self-attraction (**Supplementary Figure S3** and **S5**), we observed that the intra-arm contact pattern of chromosome 4 made of strips (~100 kb thick) was well captured only for mild interaction strength. As the AC interaction strength was increased, the separation between enriched and depleted strips became sharper with the effect of degrading the similarity of the CS with the Hi-C map.

The highest SCC value of 0.83 and the optimal similarity with the Hi-C compartment strength values were obtained for the CH-repulsive scenario (E_CH-*_~0.0004 k_B_T) together with self-attraction within active chromatin and facultative heterochromatin (E_AC-AC_=0.20 k_B_T and E_FH-FH_=0.50 k_B_T) regions (**Figures 2S-T**). Hence, this set of interactions was used to generate genome-wide models.

### Genome-wide models reveal an overall preferential V-shape for *A. thaliana* chromosomes

To generate genome-wide models, we prepared each of the 10 model chromosomes (chromosomes 1 to 5 in 2 copies each, **Supplementary Table S1**) as rod-like objects made of stacked rosettes along the main axis (29, 64), each mimicking a simplified shape of elongated mitotic chromosomes (**Supplementary Videos S2** and **Supplementary Figure S6A**). The positions and orientations of each chromosome were chosen randomly inside a sphere of diameter 5.0 micrometre (μm), that is the typical *A. thaliana* nuclear size (81).

From 50 independent replicates of these initial chromosome conformations (**Figure 3B**), we simulated a few hours of the full genome dynamics (64) applying the optimal set of parameters inferred from the single-chromosomes simulations (**Figure 3A**). To characterize the contact patterns of the obtained models (representative snapshot in **Figure 3C**), we computed the genome-wide contact map (**Figure 3D**) and the average probability of contact P(s) as a function of the genomic distance s between genomic regions on a set of ~2,000 conformations and compared them with the genome-wide interaction map and the P(s) obtained from Hi-C experiments (44, 45) (**Figure 3E** and **Material and Methods**).

From the models, we recovered the contact patterns of the Hi-C at the genome-wide scale. Cis-chromosome areas of the contact map (see black dashed squares in **Figure 3D**) had much more contacts than the trans-chromosome ones. This feature indicates that the models well captured the organization of the nucleus into distinct chromosome territories (41, 42, 44, 67). The epigenomics-driven models also recapitulated the contact enrichment between chromocenters of different chromosomes (see brown squares in **Figure 3D**) indicating that the effective repulsions between CH and the other chromatin states recover both the segregation of chromocenters at the cis-chromosome scale and the effective trans-chromosome attraction between chromocenters (41, 42, 44–46, 52, 67). To quantify the similarities of the models with experiments, we computed the CS for each epigenomic state and the SCC between the genome-wide maps, the 5 cis-chromosome maps (**Figure 3F**) in Hi-C and in our predictions. The comparisons between contact maps resulted in similar CS distributions (**Supplementary Figure S8A**) and significant SCC values. We found that the minimal SCC of 0.44 was obtained for the genome-wide contact maps, where the size of the compared samples is ~7,000,000, and that in the per-chromosome comparisons the SCCs were always larger than 0.70 (sample-size > 180,000).

Interestingly, we found a discrepancy between the experimental and the predicted contact probabilities P(s) (**Figure 3E**). The experimental P(s) exhibits two regimes: at short and intermediate genomic distances (100 kbp < s < 10 Mbp) P(s) decays as s^−0.84^, which is very well captured by the models. But, at larger genomic distances (s > 10 Mbp), corresponding to the typical range of inter-arm contacts, the observed increase of P(s) in Hi-C is missing in our prediction. Confirming the visual impression, we found a weak SCC value (=0.22 for sample-size=1,014) when comparing the predicted and observed P(s).

To account for the inter-arm increase of contacts in the Hi-C maps, we designed a novel strategy to precondition the chromosome models as V-shaped arrangements (**Figure 3G**). The preconditioning attempted to incorporate an effective memory of chromosome structure throughout the cell cycle, which was suggested by Carl Rabl already in 1885 (94) based on microscopy observations of dividing cells in salamanders. Additional elements in favour of a persistent V-shape organization of interphase chromosomes were provided by modelling studies in several species, including yeast (95–98), drosophila (99), and human (100). In *A. thaliana*, during anaphase chromosomes are pulled, centromeres first, towards opposite poles of the mother cell (52, 101). This results in a V-shape organization for metacentric chromosomes 1, 3 and 5 and in hook-like structures for acrocentric chromosomes 2 and 4. Assuming that chromosomes exhibit inherent properties of long-polymers in dense or semi-dilute solutions (31, 32, 102) we hypothesise that chromosomes will maintain an effective memory of these V-like shapes during interphase.

Accordingly, we initially arranged each chromosome in the linear (rod-like) shape (**Supplementary Figure S6A**) and then pulled it by the kinetochore (centromere) with harmonic forces along parallel directions (**Supplementary Video S3** and **Supplementary Figure S6B**). To allow for the dragging of the entire chromosome structure, during the pulling process we pinned the chromosomes in a looped conformation using harmonic bonds bridging regions at a typical separation of ~40 kbp (14 model beads) (**Supplementary Methods**).

We simulated 50 replicates of the system with V-shaped chromosomes using the optimal epigenomics-driven interactions and characterized quantitatively the obtained conformations (representative snapshots in **Figures 3G-H**) computing the genome-wide contact map and the P(s) (**Figures 3I** and **3J**). Applying the SCC analysis to compare with Hi-C, we observed that preconditioning the chromosomes in V-shaped conformations allowed capturing qualitatively and quantitatively the behaviour of the P(s), whose model *vs.* Hi-C SCC increased dramatically from 0.22 to 0.88 (**Figures 3J** and **3K**). The comparison of the contact maps (**Figure 3K**) were slightly improved (between 4 and 6% in SCC) for the genome-wide and the cis-chromosome cases of the acrocentric chromosomes 2 and 4, and the metacentric chromosome 3. SCC of the other metacentric chromosomes (chromosomes 1 and 5) were only marginally degraded (3 and 2% in SCC respectively). The distributions of compartment strength (**Supplementary Figure S8**) were shifted towards larger values in the V-shaped chromosomes, but yet they were largely consistent with the Hi-C ones. The overall improvement of the results with V-shaped chromosomes prompted us to use this chromosome shape for the rest of the genome-wide simulations.

Interestingly, we also tested the possible formation of V-shaped chromosomes from a radial chromosome pulling which is less biologically-founded (**Figures 3L-P**, **Supplementary Video S4**, **Supplementary Figure S6C**, and **Material and Methods**). We found overall lower correlations with the Hi-C (**Figure 3P**). In particular, the SCC of the P(s) dropped from 0.88 for the parallel pulling case to 0.78 for the radial one.

### The epigenomics-driven models capture the nuclear organization in *A. thaliana*

To further characterise the models obtained from the optimal set of parameters, we looked at the preferential nuclear location of the regions assigned to each epigenomic state (**Figure 4A-E**) and computed the distribution of their radial positions (**Figures 4F-J**, **Supplementary Figure S9**, and **Material and Methods**). To disentangle which of the typical nuclear positioning was to attribute to the specific epigenomics-based interactions, we designed and performed a reference set of simulations in which the initial V-shaped chromosome positioning and the NORs and telomeres attractions were maintained, but the other interactions were removed (see networks in **Supplementary Figures S7-8B**). The results of this variant system are shown in the histograms in **Figures 4F-J** in white colour as a term of comparison with the optimal interaction model whose results are shown in the characteristic colour of the epigenomic state.

Interestingly, we found that the nucleolus typically assumed a round shape with a radius around 1,250 nm and occupied the centre of the model nucleus (**Figures 4A** and **4F**). The telomeres tended to localize at the nucleolar periphery (**Figures 4B** and **4G**). These features were consistent with the reference model in which NORs and telomeres were involved in the same interactions. Notably, the constitutive heterochromatin (CH) domains typically occupied the outermost shell of the nucleus in the optimal model but not in the reference one (**Figure 4C** and **4H**), in which the CH repulsions were removed. The active-chromatin and the facultative heterochromatin also tended to a slightly more peripheral positioning than in the reference interaction model (**Figures 4D-E** and **4I-J**). Overall, these results are consistent with experimental evidence on the typical positioning of the nucleolus at the nuclear centre (41, 49, 50), of telomeres at the nucleolar periphery (42, 44–46, 67), and of heterochromatic regions at the nuclear periphery (46, 52).

Next, we tested whether the preferential locations of the nucleolus and heterochromatin are also consistent with the fact that the telomeres, and chromocenters of chromosomes 2 and 4 (which host the NORs) associate with the nucleolus forming the so-called nucleolar-associated domains (NADs) (50), which are stable landmarks of the *A. thaliana* genome organization and are maintained under heat stress conditions (103). In *A. thaliana*, NADs correspond to repetitive elements that are transcriptionally silenced by repressive histone modifications and DNA methylation. To identify the predicted NADs in the nuclear models, we computed per each 3kbp-region (1 bead) in the models the number of contacts with the NORs particles within a distance cutoff of 200 nm (**Material and Methods**). In agreement with the experiments (50), we found that the top 10% regions making contacts with the NORs particles are the telomeric regions of each chromosome and the short arms of chromosomes 2 and 4 (**Figure 4K**). As a consequence, also the chromocenters of chromosomes 2 and 4 are involved in many contacts with the nucleolus and are typically found in a perinucleolar location in agreement with experimental data (46, 52).

Next, we tested whether the typical location of the heterochromatic regions at the nuclear periphery is also consistent with the experimental evidence that groups of chromocenters coalesce together in 8.6+/-0.2 distinct foci (46). Specifically, in each snapshot of the trajectories, we considered the regions composing each of the 10 centromeres (as a proxy of the chromocenters, **Supplementary Table S1**) and associated to each of them a sphere centred at the centre of mass of the beads with a radius equal to their radius of gyration (**Figure 4L**). Per each chromocenters pair, we computed the overlap volume between the two representative spheres. To compare with experiments (46), we selected the threshold of the significant overlap to 0.34 so that the average number of distinct chromocenters in the model nuclei over the 50 replicates simulations is 8.6 that matches the average number measured experimentally (8.6+/-0.2). Overall, the distribution computed from the models is slightly skewed towards larger numbers of foci than the experimental one (**Figure 4M**). Yet, the corresponding distribution of chromocenters’ numbers resembles the experiments with all the predicted values per bin showing no significant differences with the experimental measures. The differences with the prediction might be also due to an underestimation of the linear size of the centromeric regions in the *A. thaliana* reference genome, which was used to define the length of the model chromosomes (104). Larger centromeric sequences would likely favour co-localisation in the models and skew the predicted distribution in **Figure 4M** closer to experimental data.

### NORs and heterochromatin interactions shape the nuclear organization in *A. thaliana*

To test the role of each epigenomics-based interaction, we generated models for eight variant cases in which we modified the interactions involving one chromatin state at a time. Specifically, we perturbed the optimal interaction model (**Figure 3A)** by removing the self-attraction among NORs and telomeric regions (**Figure 5A** and **Supplementary Figure S12**), removing the repulsions between CH beads and the other epigenomic states (**Figure 6A** and **Supplementary Figure S15**), removing the self-attraction between AC regions (**Supplementary Figure S10**), and removing the self-attraction between FH beads (**Supplementary Figure S11**). To further test the role of the nucleolus and of the constitutive heterochromatin, we varied the size of the NORs particles (**Figures 5B-C** and **Supplementary Figures S13-S14**), and tested the alternative CH interactions schemes of **Figures 2B** and **2C**, where the constitutive heterochromatin was self-attractive and self-attractive+repulsive respectively (**Supplementary Figures S15-S16**).

**Figure 6.**
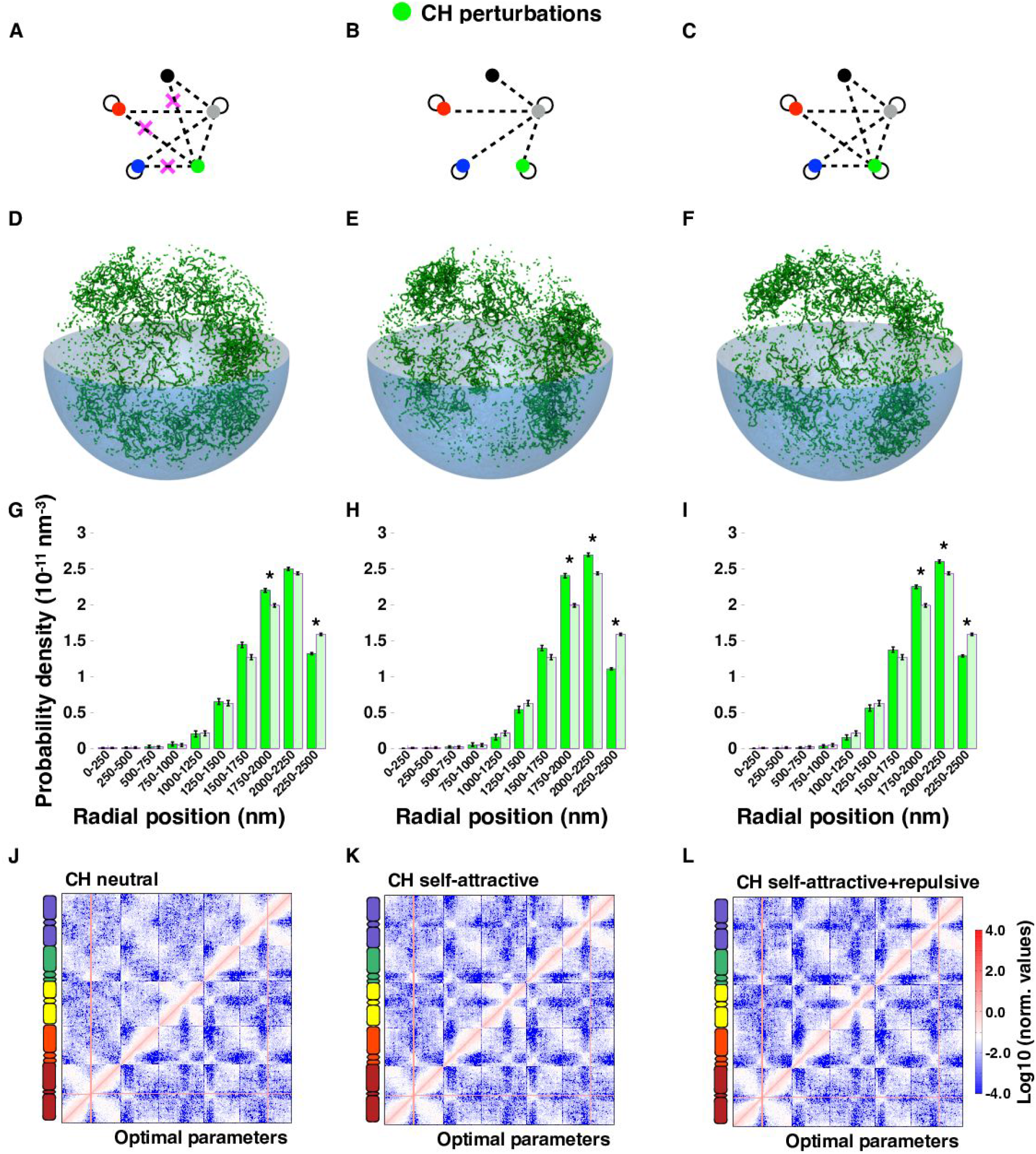
The constitutive heterochromatin interactions impact their specific nuclear positioning. **(A-C)** The variant interaction networks for constitutive heterochromatin are illustrated using the same conventions as in **Figure 2**. The interactions that are removed from the optimal case are crossed in magenta. **(D-F)** Illustrative snapshots for the CH beads are shown for the interaction models in panels scenarios in A-C. **(G-I)** Distributions of the radial positions of the 3kbp-regions in the perturbed systems (dark colour) compared to the optimal interaction model (light colour). Significant differences (two-sided Wilcoxon test p-value<0.0001) are marked with asterisks. **(J-L)** Genome-wide contact maps for the perturbed cases (top left triangles) vs maps obtained from the optimal interaction model (bottom right triangles). Complementary results on the simulations presented here are shown in **Supplementary Figures S15-S17**.

Each of the variants was compared to the optimal model by considering the radial distributions of chromatin states and by performing a SCC analysis on the genome-wide contact maps.

We found that removing only the self-attraction of the NORs and the telomeric beads and maintaining the other epigenomic-driven interactions caused dramatic nuclear rearrangements involving all the chromosomes. As expected, the compact nucleolus was disrupted (**Figure 5D**, representative snapshot) and the NORs regions were spread over the entire nucleus, resulting in an enrichment of NORs beads at the nuclear periphery and a depletion at the centre with respect to the optimal interaction model (**Figure 5G**). Interestingly, these findings are qualitatively consistent with the imaging data in human Dnmt1-deficient cells where the nucleolus is disrupted and the nucleolar rRNA genes are scattered throughout the nucleus (105). The telomeres expectedly lost their preferential perinucleolar positioning and relocalized in more peripheral shells of the nucleus (**Supplementary Figure 12B**). The perturbation of NORs interactions also affected the nuclear positioning of the constitutive heterochromatin, active chromatin and facultative heterochromatin, that are found in more central nuclear positions despite the maintenance of the epigenomics-driven interactions (**Supplementary Figures S12C-E**).

Varying the size of the self-attracting NORs particles had also large impacts on the nuclear positionings of all the chromatin states. Small NORs beads induced the formation of a smaller nucleolus of radius about 1 μm (**Figures 5E-H**) and allowed all the other chromatin states to occupy more central nuclear positions (**Supplementary Figure S13C-E**). In particular, the constitutive heterochromatin (which maintained its repulsive interactions) lost its preferential peripheral positioning (**Figure 5K**) indicating that, in our prediction, the nucleolar push is necessary to recapitulate the expected CH positioning at the nuclear periphery. This is confirmed by the simulations performed with larger NORs beads that lead to a nucleolus that occupies almost the entire nucleus (**Figures 5F** and **I**) and that pushes all the chromatin states towards the nuclear periphery with an enhanced effect for CH (**Figure 5L** and **Supplementary Figures S14C-E**).

Interestingly, the large changes in nuclear positioning induced by each of the NORs’ variant systems had a marginal effect on the respective contact patterns. The corresponding genome-wide contact maps, that did not include the NORs regions for consistency with the Hi-C interaction maps, appeared to be visually very similar to the one obtained for the optimal interaction model (**Figures 5M-O**) and quantitatively correlated equally well with the Hi-C interaction maps (**Supplementary Figures S7C-E**).

The removal of the repulsive interactions between CH beads and the other epigenomic states (**Figure 6A**) had the effect to push the heterochromatic regions towards the nuclear centre so that CH beads are less probably found in the outermost nuclear shell than in the optimal interaction model (**Figures 6D** and **6G**). Notably, the CH neutrality also marginally affected the nuclear location of the other epigenomic (AC and FH) states by pushing them slightly towards the nuclear centre (**Supplementary Figures S15D-E**). Interestingly, the main reverberation of the CH perturbation appears in the genome-wide contact map. Specifically, the signatures of the segregation of the centromeric regions are visually lost, and the trans-chromosome sections of the map are different from the correspondent parts in the optimal interaction case (**Figure 6J**). Quantitatively, the similarity of the model and the Hi-C contacts map were degraded with respect to the optimal model both in terms of the Spearman correlation (SCC=0.43 vs SCC=0.49 for the optimal), and the CS distribution that was only marginally matching the Hi-C (**Supplementary Figures S7F** and **8F**). Although the average contact probability as a function of the genomic separation, P(s), is captured accurately (SCC=0.93) (**Supplementary Figure S7F**). The two variant simulations in which the constitutive heterochromatin (CH) was involved in purely self-attractive (**Figure 6E**) and in combined self-attractive and repulsive (**Figure 6F**) interactions also showed significant differences with the optimal model case. Neither of the two scenarios leaded to a significant peripheral localization of CH as the optimal case (**Figures 6H** and **6I**) demonstrating that, although the overall compliance with the Hi-C data (**Figures 6K** and **6L**) is similar with the optimal case both in terms of Spearman correlation (**Supplementary Figures S7G-H**) and compartment strength (**Supplementary Figures 8G** and **H**), the different types of CH-interactions had an impact on its recruitment at the nuclear periphery, that could in turn precondition CH to a tethering with the nuclear lamina. Similarly to CH, AC and FH were also significantly brought towards the nuclear center in the alternative scenarios (**Supplementary Figures S15-S17**).

The perturbations of the AC and FH interactions (**Supplementary Figures S10-S11**), that involved a sizeable fraction of the genome (52% and 14%, respectively) had only negligible effects on both the radial positioning of the corresponding chromosome regions and the genome-wide contact pattern.

### The model plasticity allows accommodating fine-scale structural properties: the KNOT Engaged Elements (KEEs) and the local polycomb-like domains

To test whether adding specific sets of interactions (both epigenomics-driven or not) may help to recover structural properties at fine-scale without compromising the ones at large-scale, we studied more in detail the structural role of the KNOT engaged elements, KEEs (aka Interacting Heterochromatic Islands (IHIs)) (42, 67) and of the facultative, polycomb-like (FH) heterochromatin (**Figure 7** and **Supplementary Figures S18-S19**). In particular, we tested whether we could recover the formation of long-range interactions between the 10 KNOT Engaged Elements (aka IHIs) that had been identified in Hi-C contact maps of *A. thaliana* (42, 67). KEEs appear as strong cis- and trans-chromosome peaks in the Hi-C interaction maps. Notably, we found that the KEEs associated peaks were completely absent in the predictions of the optimal epigenomics-driven model (**Supplementary Figure S19A**) indicating that it is unlikely that such contacts can be promoted by epigenomics-driven interactions alone. In fact, in our analysis the KEEs were not enriched in any of the four epigenomic states (**Supplementary Figure S18**), consistent with other ChIP-seq analysis that suggested that KEEs are not associated with a specific chromatin state (42, 44, 60).

**Figure 7.**
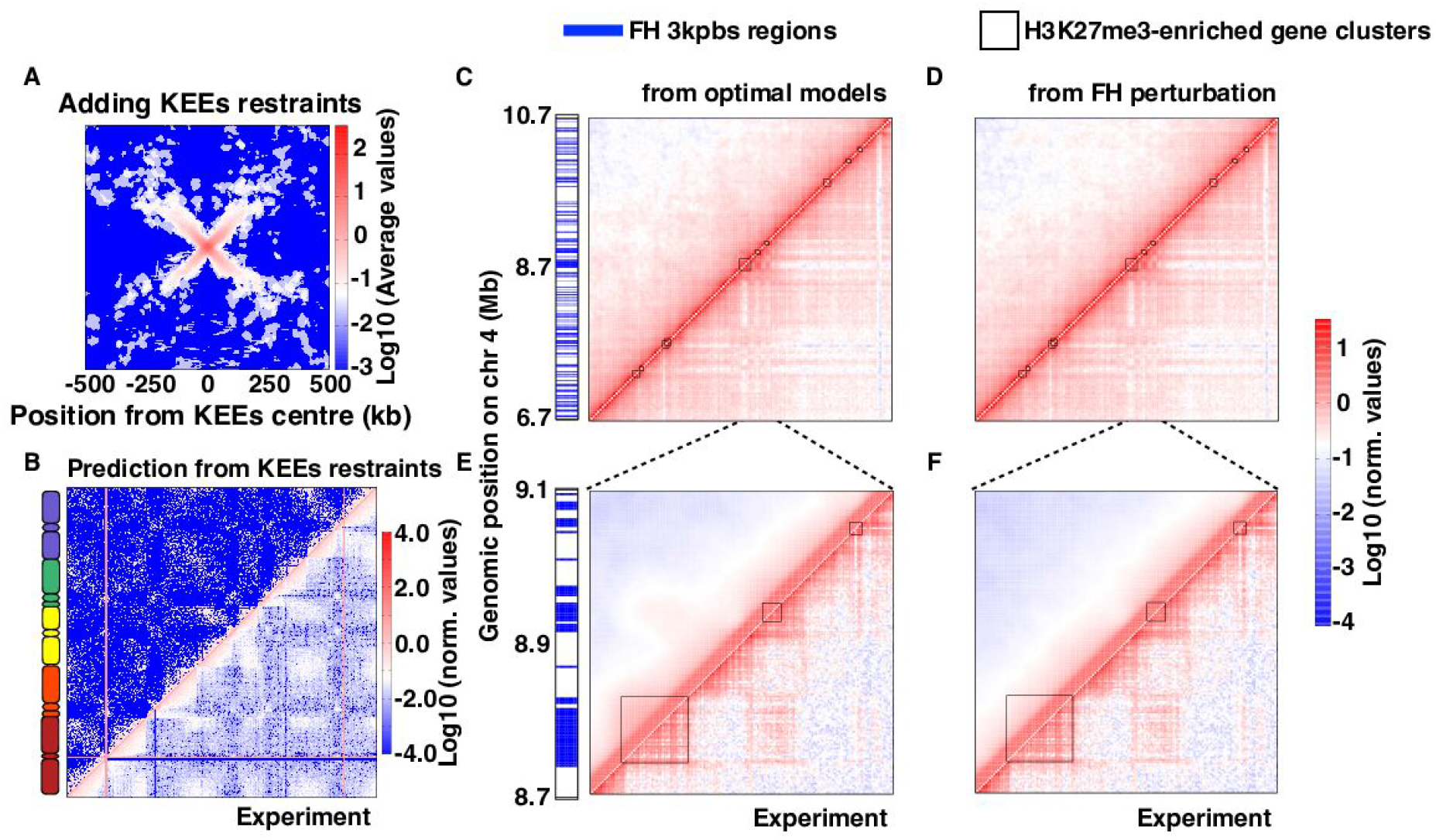
Plasticity of the optimal models in response to long-range and local interactions. **(A)** Average contact map of a 1Mbp-region around the KNOT Engaged Elements (KEEs) for the KEEs-restrained models. **(B)** Genome-wide contact maps in the KEEs-restrained case (top left triangle) and from experiments (bottom right triangle). **(C-D)** Predicted (top left triangles) vs experimental (bottom right triangles) contact maps at 30 kbp resolution in the chr4:6,700,000-10,700,000 bp region for the optimal set of parameters used in **Figures 3** and **4** (**C**) or the FH-perturbed case in which the self-attraction between polycomb-like beads is switched off (**D**). **(E-F)** Zoomed views of **(C-D)** around the largest gene cluster (cluster 100 of Ref. (56)) (chr4:8,668,000-9,109,000 bp at 3 kb resolution). Black squares show the positions and extents of the gene clusters enriched in H3K27me3 mark (56). The formation of polycomb-promoted domains, which is recapitulated in the optimal model (**E**), fades away in the absence of FH self-attraction (**F**). On the left of each map, blue lines indicate the positions of polycomb-like beads.

Association rates of 4 KEEs’ pairs (KEE6-KEE1 20%, KEE5-KEE4 35%, KEE6-KEE3 66%, KEE5-KEE10 16% with an average of 34%) were previously measured by Fluorescent In Situ Hybridization (FISH) (67), which showed an average of 34% co-localization of KEEs pair. To enforce their mutual spatial proximity, we applied long-range harmonics (both cis- and trans-chromosomes) between the central beads of KEEs’ pairs (**Material and Methods**). To use the minimal set of harmonics and the minimal possible forces, we computed the distances between all the pairs of KEEs central beads (both in cis- and trans-chromosome) in the 50 final snapshots obtained from the optimal model simulations and ranked the pairs for increasing mutual distance. On the same 50 snapshots (representative snapshot in **Figure 3H**), we applied harmonics (**Material and Methods**) between the closest 34% (or, when available, the specific FISH association rate (67)) of each KEEs pair to promote their spatial colocalization.

We found that the plasticity of the models allowed accommodating all the KEEs interactions and that the generated contact pattern around KEEs pairs was a cross appearing in the inter KEEs portions of the contact map (**Figure 7A**). This specific pattern was due to the applied constraints that bring into spatial proximity only the central beads. However, recovering the KEEs contacts largely affected the large-scale chromosome rearrangements as it is conveyed visually by the genome-wide contact map (**Figure 7B**) and quantitatively by the Spearman correlation with the Hi-C interaction map, that degrades both genome-wide and in each cis-chromosome parts with respect to the optimal interaction model (**Supplementary Figure S19D**).

Next, we focussed on local structural features involving the 162 gene clusters enriched in the H3K27me3 polycomb-related histone mark (median length 7.5 kbp and maximum length 95 kbp) which have been identified in the *A. thaliana* genome (56). In the Hi-C interaction map, these clusters form a local plaid pattern indicating the presence of interactions within and between them. We considered a 4 Mbp region (chr4:6,700,000-10,700,000) (**Figure 7C**) centred at the longest gene cluster, the 95kb-long cluster-100 of Ref. (56). Interestingly, we observed that the FH self-attraction already present in the optimal models (**Figure 3A**) were primed to capture the formation of these local contact domains between H3K27me3-enriched gene clusters (**Figure 7E**). These domains were not recapitulated in the perturbed system where the FH self-attraction is removed (**Figures 7D-F**). This finding suggests that we could recover structural features at the scale of a few hundreds of kilobases, that were close to the limit of the model coarse-graining (~300 kb).

## DISCUSSION AND CONCLUSION

We studied how and to what extent epigenomics-driven interactions shape the structural organization of the *Arabidopsis thaliana* genome. For this purpose, we developed a computational strategy based on an underlying polymer model that, decorated with epigenomics-based interactions (attractions and repulsions) and simulated via molecular dynamics, allowed generating accurate genome-wide structures of the *A. thaliana* chromosomes coarse-grained at 3 kbp.

Overall, we found a constitutive organizational role for the attraction between the nucleolar organizing regions NOR2 and NOR4, and for the repulsions of the constitutive heterochromatin (CH) regions and the rest of the genome. Interestingly, these interactions, that are part of the optimal interaction model, allow recovering several established experimental results (**Figure 3** and **4**). Specifically, the formation and the central nuclear positioning of the nucleolus (41, 49, 50), the close positioning of NADs to the nucleolus itself (50, 103), the peripheral positioning of the CH regions (46, 52, 106), and the coalescence of chromocenters in discrete *foci* (42, 46, 67) were all recapitulated in our models. Importantly, these accurate predictions were lost in alternative models we tested. When we removed the NORs self-attraction, or varied the NORs particles size, the positioning of all epigenomic domains were strongly perturbed (**Figure 5** and **Supplementary Figures S12-S14**). These tests indicate that the nucleolus has a major effect on the nuclear organization including the peripheral location of constitutive heterochromatin. When we removed the CH repulsions or we substituted it with CH self-attraction, the peripheral positioning of heterochromatic regions was also lost (**Figure 6** and **Supplementary Figures S15-S16**).

The major role of nucleolus and centromeres to shape the nuclear organization in *A. thaliana* is consistent with previous findings from genome modelling (58), that we largely extend by providing mechanistic insight. We modelled the formation of the nucleolus by the aggregation of the self-attractive constitutive NORs regions and by preconditioning them at the nuclear centre. We also tested that the NOR self-attractions are essential to maintain both a compact and central nucleolus (**Figure 5**) implying that the preconditioning and non-specific interactions, such as the depletion effect which may act due to the different size of the NOR beads, are not enough to achieve accurate structural organization. Our computational protocol for the formation of the nucleolus is also one of the first attempts of its kind in the modelling of the 3D genome organization higher eukaryotes. The only exception is yeast in which the rDNA sequence on chromosome XII has been modelled with an effective repulsion (96) or with *ad hoc* confinements (97, 98).

Our findings also reveal that the segregation of the chromocenters, their preferential positioning at the nuclear periphery, and their association in discrete clusters can be explained by epigenomics-driven effective repulsions of the constitutive heterochromatin with the other epigenomic states. In *A. thaliana*, the nuclear envelope hosts the nucleoskeleton (107), a peripheral matrix that is functionally similar to the nuclear lamina in animal cells (108, 109). It has been demonstrated that the proteins *crwn*1 in presence of *crwn4* and non-CG DNA methylation (107, 110) can bridge to the nucleoskeleton specific genomic regions, called Plant Lamina-Associated Domains (PLADs). Similarly to the LADs in animals (74, 111, 112), PLADs are enriched in repressive chromatin marks and silenced chromatin but are not in perfect correspondence with heterochromatin. Here, our simulations suggest that the repulsion of the CH with the other chromatin states may favour the recruitment of chromocenters at the nuclear periphery significantly more than the CH self-attraction (**Figure 6H**) and precondition these regions for the dynamical tethering at the peripheral matrix by *crwn1* and *crwn4* proteins. However, it is also likely that the CH self-attraction scenario combined with attractive interactions between PLADs and the nucleoskeleton may also lead to a proper peripheral positioning as suggested by recent models in mammals (74, 112).

Together with NORs and CH interactions, another important finding of our study is that different initial shapes of chromosomes have a huge impact in recovering experimental evidence. We showed here that preconditioning the chromosome structures in a V-shape state, as expected for (sub)metacentric chromosomes, dramatically improves the description of the contact probability vs. the genomic distance, P(s) (**Figures 3I-J**). This finding is to ascribe to topological constraints that allow long polymers (that are viable models for chromosomes (113)) keeping a partial memory of their initial arrangements over long timescales (29, 32, 102). The models allow discerning that the V-shapes obtained from the parallel (not radial) pulling of all chromosomes by the centromeres generate the most accurate models (**Figures 3K** and **3P**). The role of the initial conformation in our simulations is emphasized by the fact that our models avoid chain crossings. In the nucleus, DNA topoisomerase type II allows DNA filaments to pass through each other, but its role in regulating the large scale chromosome organization is still highly unclear. It has been suggested, in fact, that facilitated chain crossing would prevent the formation of chromosome territories (114), lead to different scaling properties for P(s) than experimentally determined (16, 115), and favour the formation of complex physical knots that is incompatible with the low entanglement observed and predicted in chromosomes (36, 116–118). However, the role of chain crossings and genome topology still deserves attention (102) since it has been shown that knots do form in eukaryotic minichromosomes *in vivo* (119) and may be favoured by the accumulation of entanglements, for instance during transcription (120).

Interestingly, our optimal parallel pulling protocol resembles to some extent the large-scale dynamics during cell division in *A. thaliana* when the chromosomes of each daughter cell, after the first transient dynamics, move in parallel, centromere first, towards opposite poles of the mother cell (52, 101). We also wish to point out that we gathered this insight by analysing the P(s) from the Hi-C datasets. This fact is a clear example that the Hi-C contact patterns, once they are properly interpreted, can inform on the large scale chromosomes organization and that 3D modelling approaches offer a viable bridge to contextualize and quantitatively reconcile imaging and Hi-C results (72, 121). This partial memory of the initial V-shape parallels the evidence collected in other species with (sub)metacentric chromosomes like in yeast, human and fly where the contact probability also exhibits an increase at genomic scales corresponding to inter-arm contacts (17, 96–100, 122).

Interestingly, the optimal interaction model we found is largely consistent with known protein- and RNA-mediated interactions which may shape the chromatin structure *in vivo*. Specifically, the formation of the nucleolus by the attraction of the NORs in chromosome 2 and 4 may suggest an architectural role for ribosomal RNA (rRNA), RNA Polymerase I, and transcription factors that are enriched at the nucleolus (123–126). The repulsion of constitutive heterochromatin (CH) is consistent with the evidence that the heterochromatin complex 1 (HP1) binds to repressive histone marks (e.g. H3K9me2) and possibly forms insoluble protein droplets (84, 85) that effectively increase the local chromatin volume. Similarly, the self-attraction of active chromatin (AC) could take effectively into account the binding of RNA PolII or transcription factors to active genes priming a micro-phase separation phenomenon (88, 89), and the self-attraction of facultative heterochromatin (FH) may model the phase-separation effects induced by polycomb group (PcG) proteins binding to repressive histone marks (90, 91).

Interestingly, comparably weak epigenomics-driven interactions have been found in similar studies on flies (37) and humans (34–36, 75–77). In the latter, the chromatin states are linearly organized into large (~100 kbp) blocks (127, 128) that concur to the formation of chromatin compartments in humans and TADs in flies. However, in *A. thaliana* the linear organization of the epigenomic states (**Figure 1A** and **Supplementary Figure S1**) (44, 59, 60) is characterized by small domains (median ~10kb) interspersed along all chromosomes, that may explain why we do not observe strong epigenomics-related compartments or TADs in *A. thaliana* (42, 44, 45, 67).

Additionally, the optimal models presented a degree of plasticity that allowed enforcing and satisfying the long-range contacts between pairs of KNOT Engaged Elements (KEEs). This mechanism might result from the molecular action of nuclear myosins that have been shown to promote long-range genomic contacts in mammals (129–131). However, we found in these simulations overall lower correlations between model contact patterns and Hi-C (**Figure 6**). This result suggests that the formation of KEEs interactions cannot be modelled by long-range harmonics applied on the optimal models, but has to be ascribed to alternative mechanisms. We may speculate, for instance, possible tethering of all (or part of) the KEEs to nuclear landmarks, as it is the case of the centromeres in yeast that are attached to the spindle pole body via microtubules (132). Also, short-range interactions could promote KEEs contacts, but, to be properly described using polymer modelling, they need a specific preconditioning procedure, such as the ones applied in this work for the nucleolus formation.

In conclusion, we show that using polymer models and epigenomics-driven interactions it is possible to predict the genome organization of *A. thaliana* and to discern the crucial role of NORs and heterochromatin in shaping its 3D genome. Additionally, we demonstrate that within the same modelling framework fine-scale genomic features, such as H3K27me3-enriched gene clusters and KEEs, can be quantitatively tested. Our fine-grained analysis unveils that our approach can also be used to test reliably and quantitatively local genomic structural features (close to the coarse-graining limit), and also demonstrates that polymer modelling can robustly propose or rule out possible mechanisms underlying the formation of the contact patterns by verifying simultaneously their effect at large and local genomic scales. The computational modelling introduced here for *A. thaliana* will help to unravel the mechanisms behind the genomic organization not only in other complex plant species such as wheat and rice but also in many eukaryotes in which the nucleolus and heterochromatin are highly conserved elements.

## DATA AVAILABILITY

The Hi-C datasets analysed during the current study are available in the Sequence Read Archive (SRA) repository with accession numbers SRR1029605 from Ref. (44), SRR2626429, SRR2626163 from Ref. (45). Epigenomic data were extracted from the Supplementary Table 4 of Ref. (44). All LAMMPS, Bash, R and Python in-house scripts for simulations and data analysis used for this study will be available upon publication.

## ACKNOWLEDGEMENTS

We are grateful to the members of the Marti-Renom and Jost labs, and to Paula Soler-Vila for useful discussions.

