## Supplementary_data for "Polymer modelling unveils the roles of heterochromatin and nucleolar organizing regions in shaping 3D genome organization in Arabidopsis thaliana"

Di Stefano *et al.*

### SUPPLEMENTARY TABLES

**Properties of the *A. thaliana* genome per chromosome.**

| Chr | Copy | Golden Path Length | NORs | Centromeres | Total Length | |
|---|---|---|---|---|---|---|
| | | base-pairs (bps) | | | base-pairs | beads |
| 1 | 1 | 30,427,671 | - | 13,700,000-15,900,000 | 30,427,671 | 10,143 |
| 1 | 2 | 30,427,671 | - | 13,700,000-15,900,000 | 30,427,671 | 10,143 |
| 2 | 1 | 19,698,289 | 1-3,600,000 | 2,450,000-5,500,000 | 23,298,289 | 7,767 |
| 2 | 2 | 19,698,289 | 1-3,600,000 | 2,450,000-5,500,000 | 23,298,289 | 7,767 |
| 3 | 1 | 23,459,830 | - | 11,300,000-14,300,000 | 23,459,830 | 7,820 |
| 3 | 2 | 23,459,830 | - | 11,300,000-14,300,000 | 23,459,830 | 7,820 |
| 4 | 1 | 18,585,056 | 1-4,000,000 | 1,800,000-5,150,000 | 22,585,056 | 7,529 |
| 4 | 2 | 18,585,056 | 1-4,000,000 | 1,800,000-5,150,000 | 22,585,056 | 7,529 |
| 5 | 1 | 26,975,502 | - | 11,000,000-13,300,000 | 26,975,502 | 8,992 |
| 5 | 2 | 26,975,502 | - | 11,000,000-13,300,000 | 26,975,502 | 8,992 |
| **All** | **2** | **238,292,696** | **-** | **-** | **253,492,696** | **84,502** |

**Supplementary Table S1**. The Golden Path Length was taken from The Arabidopsis Information Resource (TAIR), ftp://ftp.arabidopsis.org/home/tair/Sequences/whole_chromosomes/, on www.arabidopsis.org, Nov 2014 [1]. The centromere positions from [2] and the NORs positions and lengths were gathered from [3].

***A. thaliana* Hi-C datasets used in this study**

| Dataset | Cell type | Restriction enzyme | ID | Read length (nt) | Valid reads |
|---|---|---|---|---|---|
| Wang2014 | Col-0 | DpnII | SRR1029605 | 200 | 73,365,540 |
| Liu2016_r1 | Col-0 | DpnII | SRR2626429 | 200 | 30,096,310 |
| Liu2016_r2 | Col-0 | DpnII | SRR2626163 | 200 | 33,210,242 |
| **Total** | **Col-0** | **DpnII** | **-** | **-** | **136,672,092** |

**Supplementary Table S2**. Taken from references [2] and [4] along with the cell type, the restriction enzyme used, the NCBI accession numbers, and the number of the valid reads retrieved after filtering using TADbit [5].

**SUPPLEMENTARY FIGURES**

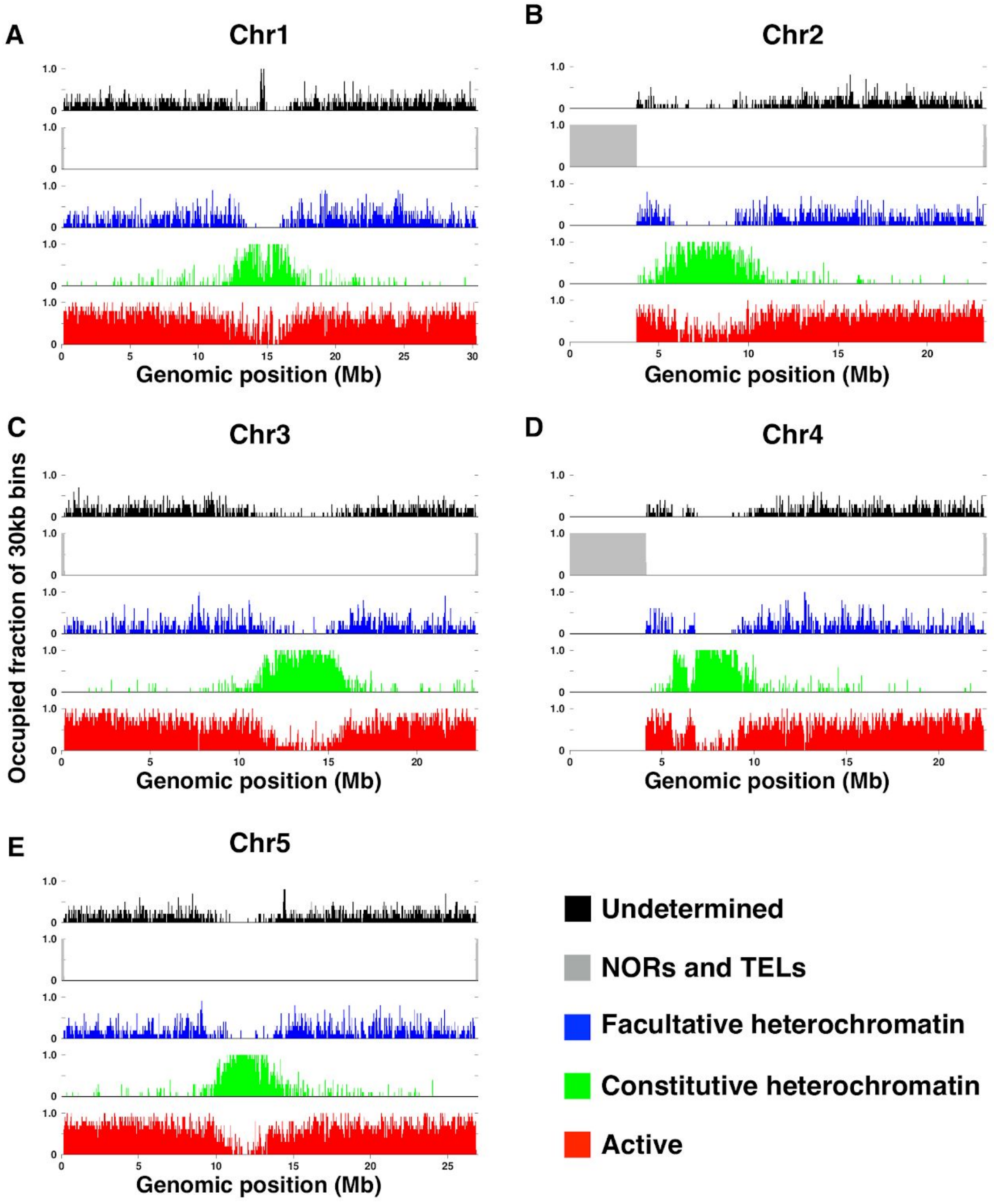


**Supplementary Figure S1. Epigenomic partition of the *A. thaliana* chromosomes.** Per each chromosome, five different bar-plots are shown, one per each of the epigenomic states. Each bar-plot shows for each 30kbp-region of chromatin the fraction (from 0 to 1) that is occupied by 3kbp-regions of the corresponding chromatin state.

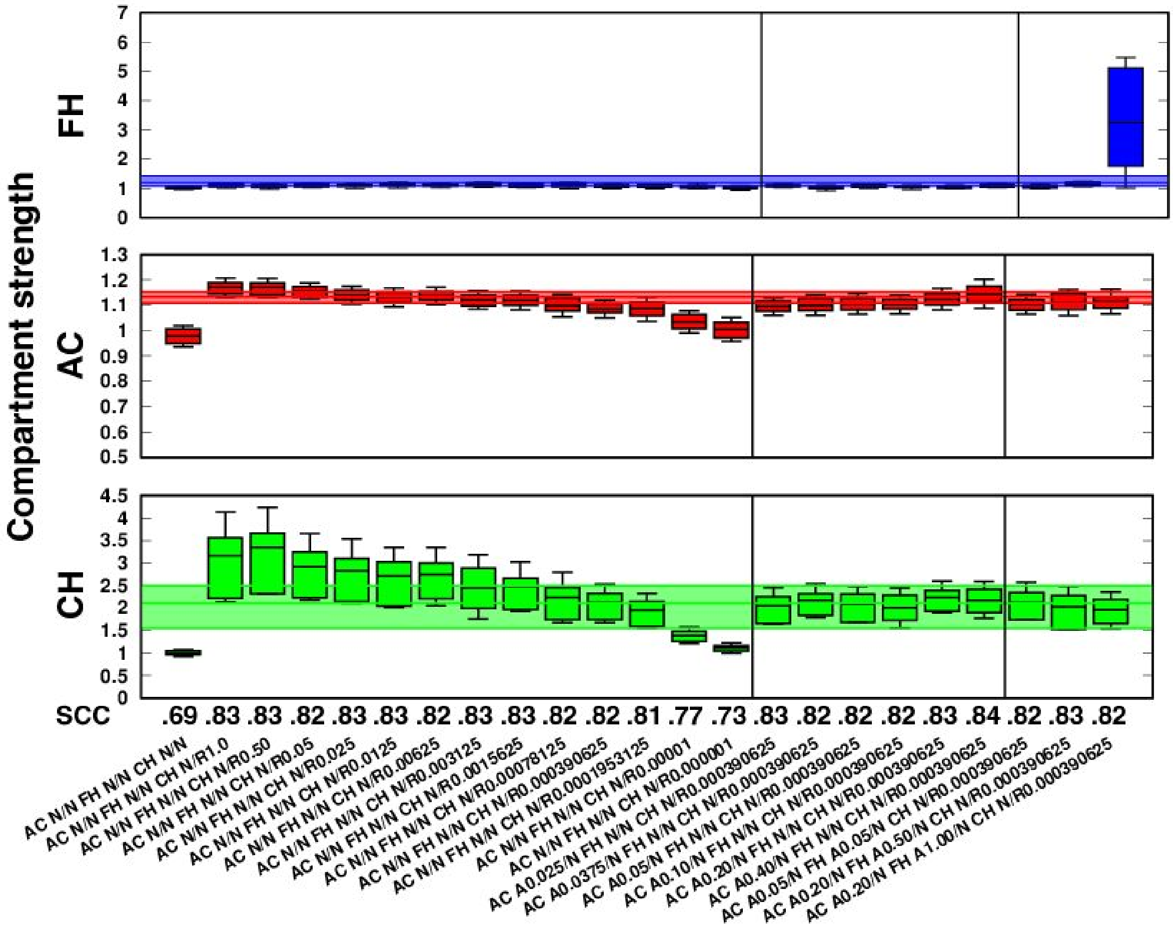


**Supplementary Figure S2. Parametrization of the single-chromosome simulations via Hi-C comparative analysis for the purely repulsive constitutive heterochromatin models.** Per each explored parameters' set simulated in 10 independent replicates, contacts in the models were counted for each pair of 3kbp-regions (beads) closer than 200 nm in space, and used to compute the contact matrix at 30kpb resolution. The models' contact maps were used to compute the Spearman correlation coefficient (SCC) with the Hi-C interaction map, and the compartment strength (CS) per bin for every epigenomic state (constitutive heterochromatin - CH, active chromatin - AC, and facultative heterochromatin - FH), which was compared with the corresponding Hi-C quantity. CS and SCC were used to select the best match between models and experimental (Hi-C) data. In this scenario, the resulting optimal model included the following interactions strengths: $E_{CH}^{rep} = 0.000390625\, k_B T$, $E_{AC}^{attr} = 0.20\, k_B T$, and $E_{FH}^{attr} = 0.50\, k_B T$. Each Hi-C CS distribution is represented in the background of the plot with a colored band which spans the range of values from the first to the third quartiles.

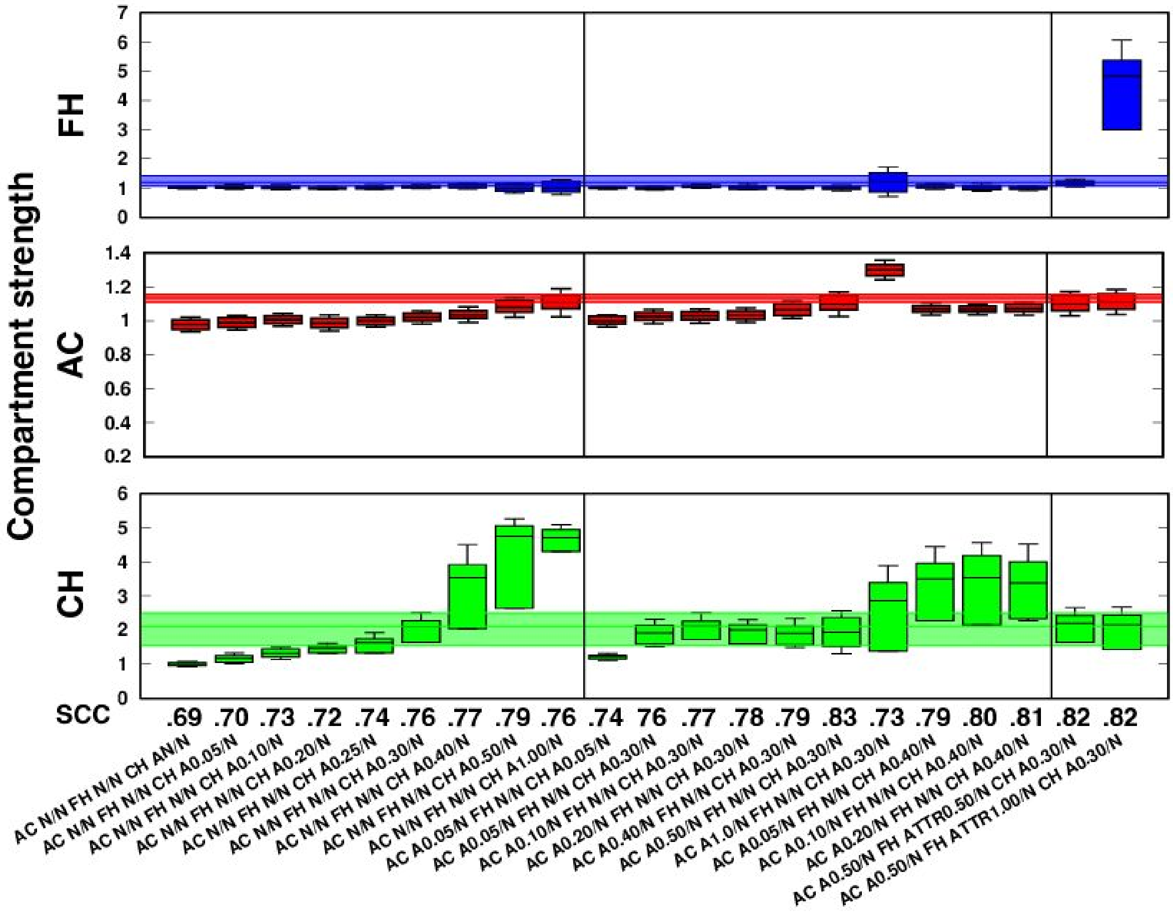


**Supplementary Figure S3. Parametrization of the single-chromosome simulations via Hi-C comparative analysis for the purely self-attractive constitutive heterochromatin models.** Per each explored parameters' set simulated in 10 independent replicates, contacts in the models were counted for each pair of 3kbp-regions (beads) closer than 200 nm in space, and used to compute the contact matrix at 30kpb resolution. The models' contact maps were used to compute the Spearman correlation coefficient (SCC) with the Hi-C interaction map, and the compartment strength (CS) per bin for every epigenomic state (constitutive heterochromatin - CH, active chromatin - AC, and facultative heterochromatin - FH), which was compared with the corresponding Hi-C quantity. CS and SCC were used to select the best match between models and experimental (Hi-C) data. In this scenario, the resulting optimal model included the following interactions strengths: $E_{CH}^{attr} = 0.30\,k_BT$, $E_{AC}^{attr} = 0.50\,k_BT$, and $E_{FH}^{attr} = 0.50\,k_BT$. Each Hi-C CS distribution is represented in the background of the plot with a colored band which spans the range of values from the first to the third quartiles.

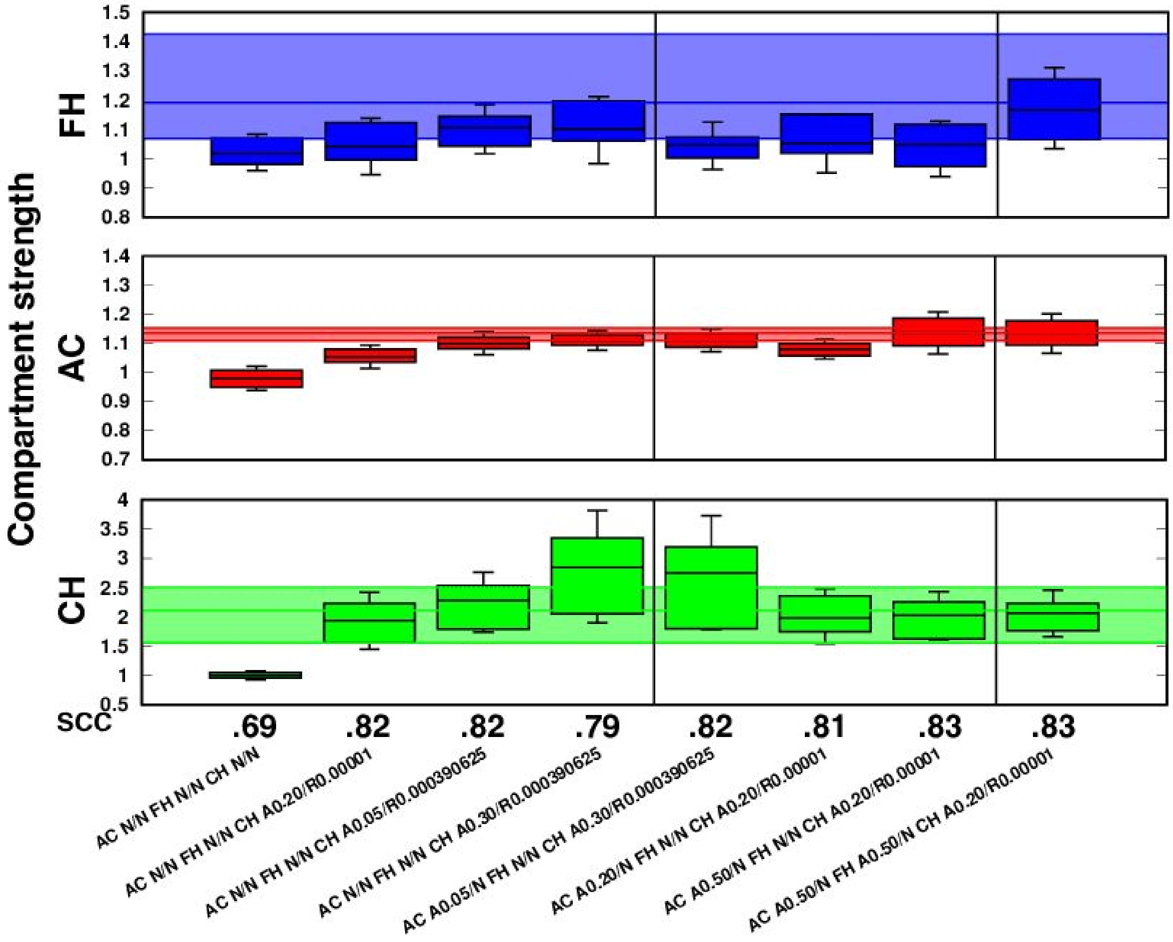


**Supplementary Figure S4. Parametrization of the single-chromosome simulations via Hi-C comparative analysis for the self-attractive and repulsive constitutive heterochromatin models.** Per each explored parameters' set simulated in 10 independent replicates, contacts in the models were counted for each pair of 3kbp-regions (beads) closer than 200 nm in space, and used to compute the contact matrix at 30kpb resolution. The models' contact maps were used to compute the Spearman correlation coefficient (SCC) with the Hi-C interaction map, and the compartment strength (CS) per bin for every epigenomic state (constitutive heterochromatin - CH, active chromatin - AC, and facultative heterochromatin - FH), which was compared with the corresponding Hi-C quantity. CS and SCC were used to select the best match between models and experimental (Hi-C) data. In this scenario, the resulting optimal model included the following interactions strengths: $E_{CH}^{rep} = 0.00001\ \mathrm{k_BT}$, $E_{CH}^{attr} = 0.20\ \mathrm{k_BT}$, $E_{AC}^{attr} = 0.50\ \mathrm{k_BT}$, and $E_{FH}^{attr} = 0.50\ \mathrm{k_BT}$. Each Hi-C CS distribution is represented in the background of the plot with a colored band which spans the range of values from the first to the third quartiles.

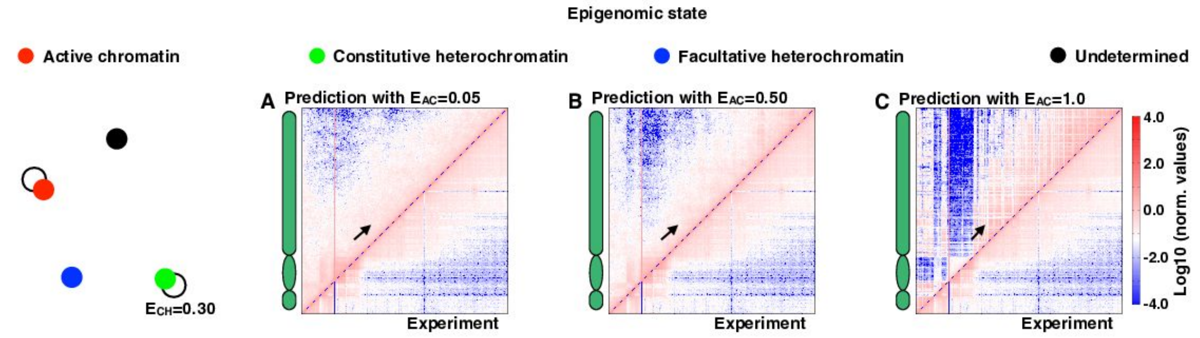


**Supplementary Figure S5. Single-chromosome contact maps at increasing AC attractive interaction strengths in the purely self-attractive constitutive heterochromatin model. (A-C)** Cis-chromosomes contact matrices of Chr4 at 30kbp resolution are shown for three increasing strengths of AC self-attraction for the interaction network model with CH and AC self-attractive interactions represented as circles in the interaction network (*Left*). At constant CH self-attraction ($E_{CH}$=0.30 $k_BT$), the increasing AC interaction strength affects the intra-arm contact patterns of models' contact maps. The mild 100kbp-thick strips of increased interactions (black arrows) are captured in the models with mild ($E_{AC}$=0.05 $k_BT$) and intermediate ($E_{AC}$=0.50 $k_BT$) AC attractions (**A-B**). But as the AC interaction strength increases further ($E_{AC}$=1.00 $k_BT$), the separation between enriched and depleted strips becomes sharper with the effect of degrading both the correlation with the contact map and the similarity of the CS distribution of the Hi-C (**Supplementary Figure S3**).

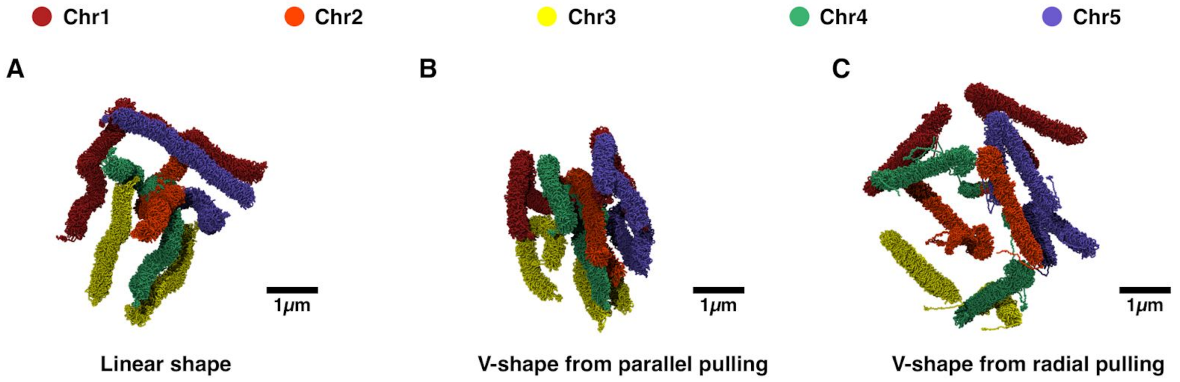


**Supplementary Figure S6. The initial shapes of the mitotic-like chromosomes. (A-C)** The genome-wide snapshots show the chromosomes at the beginning of the simulations for the different tested mitotic-like initial arrangements: **(A)** the linear shape, **(B)** the V-shape from parallel chromosome pulling, and **(C)** the V-shape from radial chromosome pulling.

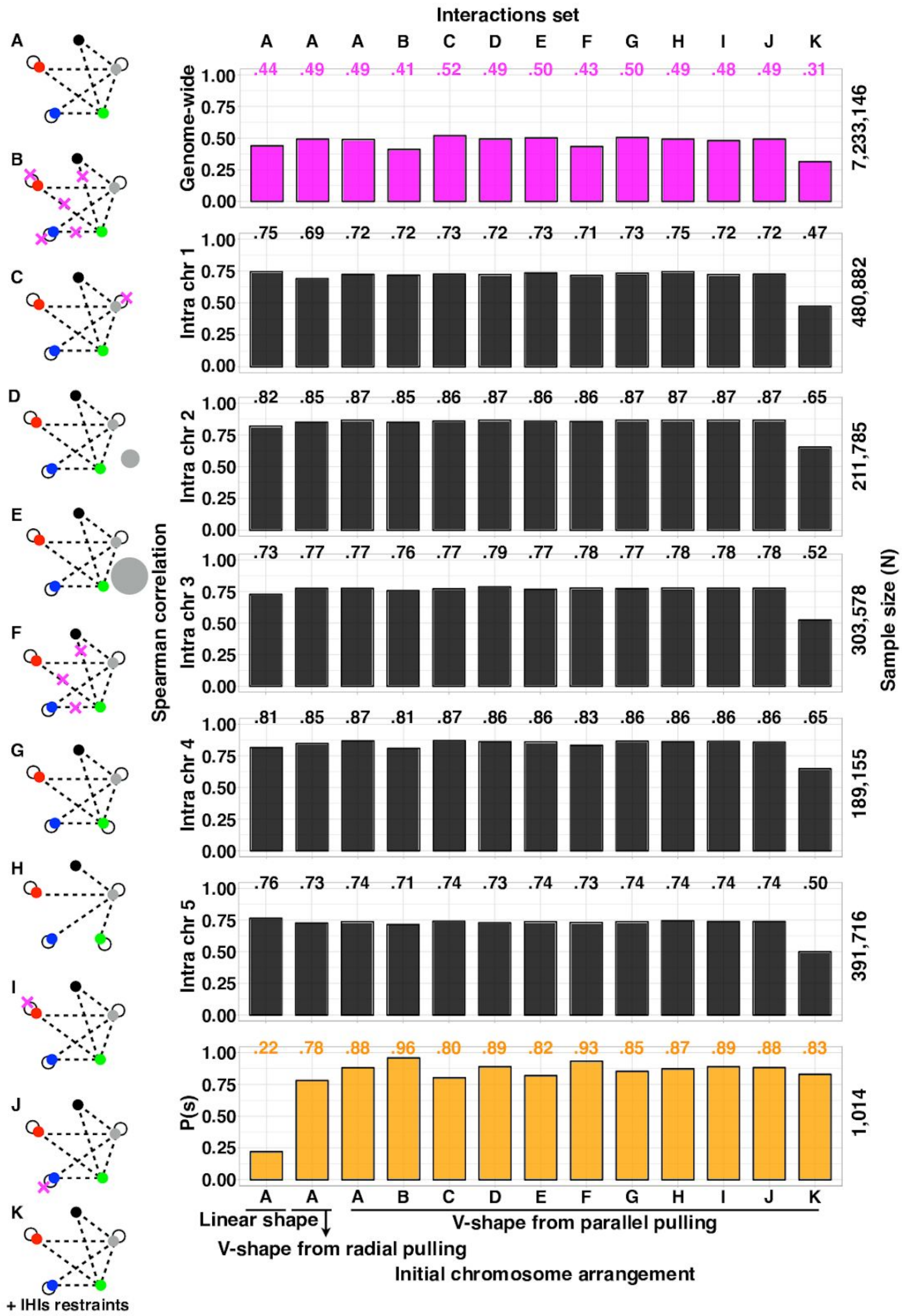


**Supplementary Figure S7**. **The account of all the explored epigenomics-driven interaction models in the genome-wide simulations and their SCC with Hi-C data.** In the genome wide simulations each interaction network (**A-K**) was explored with 50 replicate runs each. In each network the points represent the five epigenomic states: NORs in gray, CH in green, FH in blue, AC in red, and UND in black. The circles and dashed lines represent self-attractions and repulsions respectively. Unless specified, the model beads interact with an excluded volume potential that allows the fibre to maintain a thickness of 30 nm and to avoid chain crossing. The optimal case of panel **A** includes the three distinct initial mitotic-like conformations of **Figure 3** and **Supplementary Figure S6**. In the perturbed systems, the magenta crosses mark removed interactions. Small (**D**) and large (**E**) gray circles represent the perturbed NORs/TELs bead diameters. (*Left*). The SCC values obtained comparing Hi-C and simulations are shown as barplots (*Right*) for genome-wide contact maps (*Genome-wide*), *intra*-chromosome contact maps (*Intra chr 1-5*), and contact probability vs. genomic distance (*P(s)*).

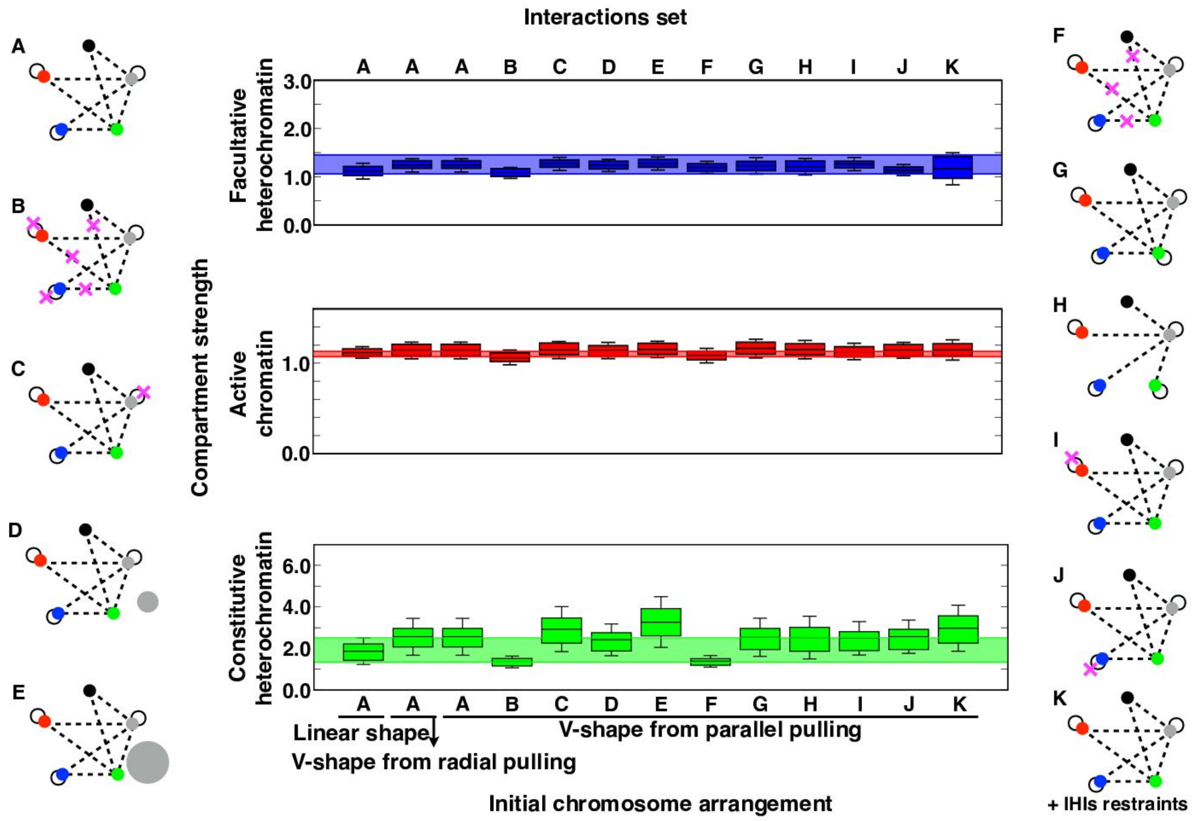


**Supplementary Figure S8**. **The account of all the explored epigenomics-driven interaction models in the genome-wide simulations and their compartment strength (CS) distributions.** In the genome wide simulations each interaction network (**A-K**) is represented as in **Supplementary Figure S7** (*Left* and *Right*). Each CS distribution from the models is represented as a boxplot, and from the Hi-C as a colored band in the background, which spans the range of values from the first to the third quartiles (*Centre*).

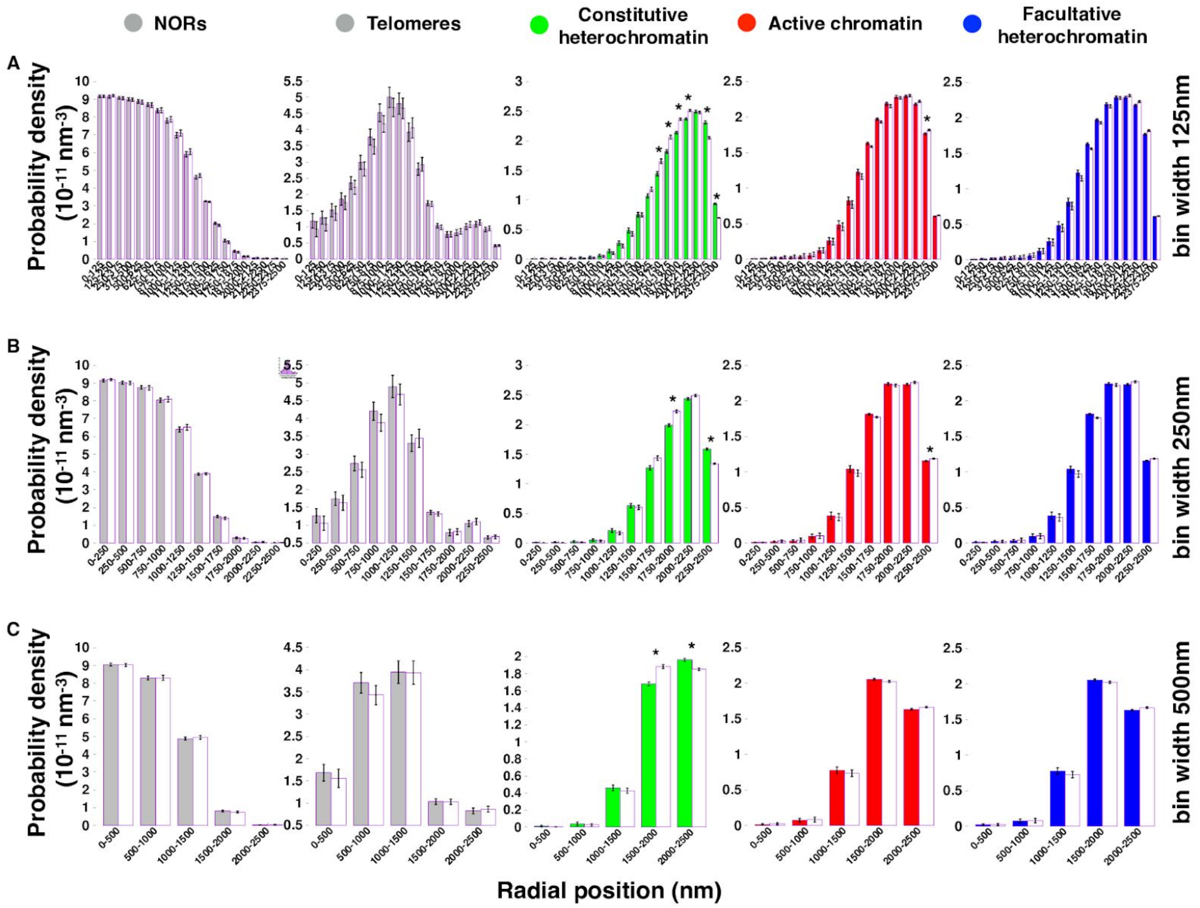


**Supplementary Figure S9. Radial positioning analysis of the epigenomic states shown in Figure 4F-J for three different radial binnings.** Bin-widths of 125 nm (**A**), 250 nm (**B**), and 500 nm (**C**) have been explored, and all show highly-consistent significant (two-sided Wilcoxon test p-value<0.0001) results. The probabilities per shell were obtained by dividing the number of particles by the volume of each shell. Histograms were computed per each replicate (1 to 50) and the means and standard errors over these replicates are reported as bar plots and error bars respectively. Wilcoxon tests are performed by comparing the distributions of the occupancies of each bin over the 50 replicates. The Wilcoxon test is suitable to compare each paired (corresponding replicates) distributions without assuming any specific shape for the two distributions under comparison. The null hypothesis is that the mean heights over the 50 replicates gave zero difference and the alternative is that the difference of the means is either higher or lower than zero (two-sided statistical test). A very stringent threshold for significance (p-value<0.0001) is chosen to single out only the most relevant differences in bins occupancies.

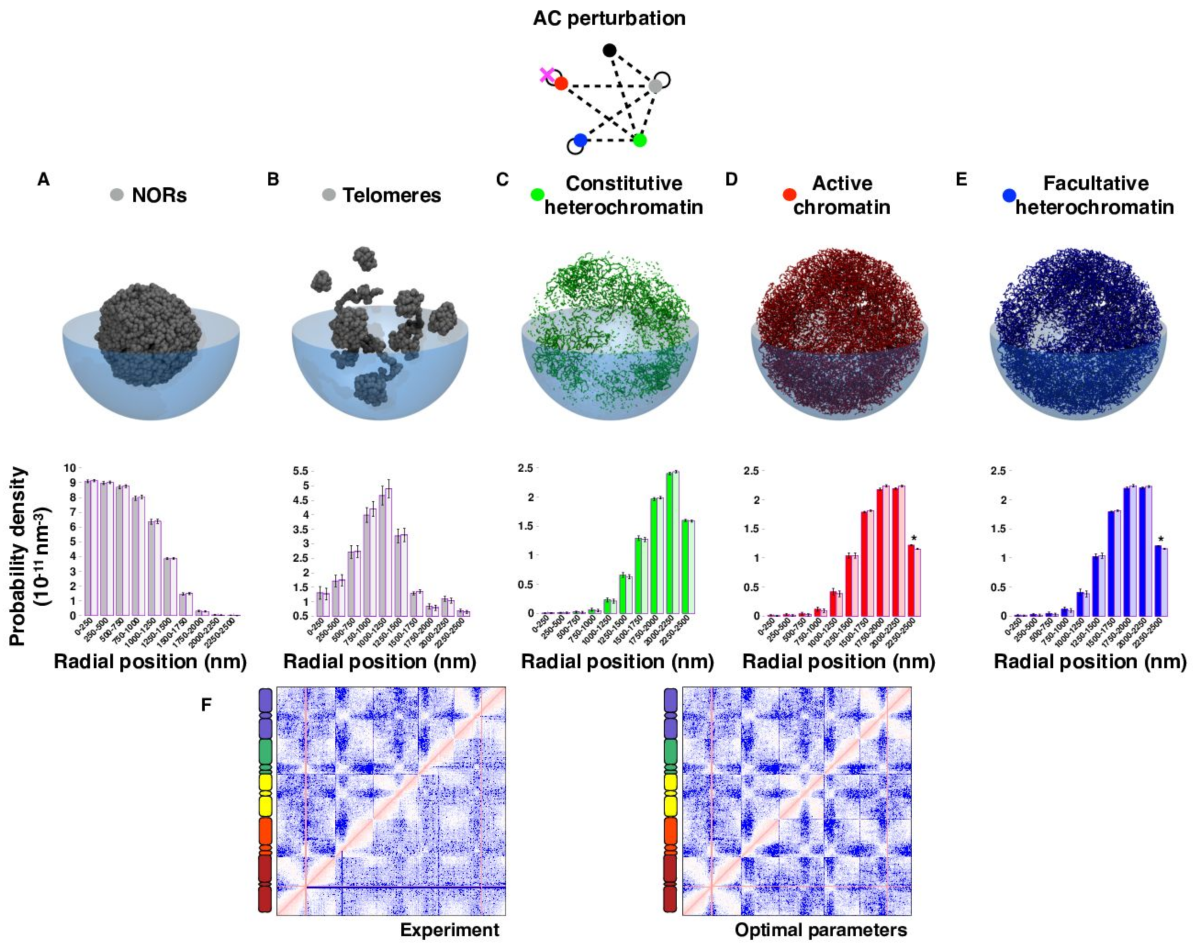


**Supplementary Figure S10. Mild effect of the removal of the self-attraction between active chromatin (AC) regions.** In the optimal interaction network the AC self-attraction was removed (magenta cross). **(A-E)** (*Top*) Each representative snapshot of the model nucleus shows the 3kbp-regions (beads) of the corresponding chromatin state in the perturbed system at the last frame of one of the 50 simulated trajectory (replicate 1). (*Bottom*) The distribution of the radial positions of the 3kbp-regions in the perturbed systems (dark colour) is compared to the optimal interaction model (light colour) of **Figure 3A**. Few significant differences (two-sided Wilcoxon test p-value<0.0001) are detected. **(F)** The genome-wide contact map for the perturbed case (top left triangles) is shown together with (*Left*) the Hi-C and (*Right*) the prediction from the optimal model (bottom right triangles). Removing AC self-attraction does not affect significantly the predicted genome wide maps and the similarity with Hi-C is almost as equivalent as for the optimal model (**Supplementary Figure S7I** and **S8I**).

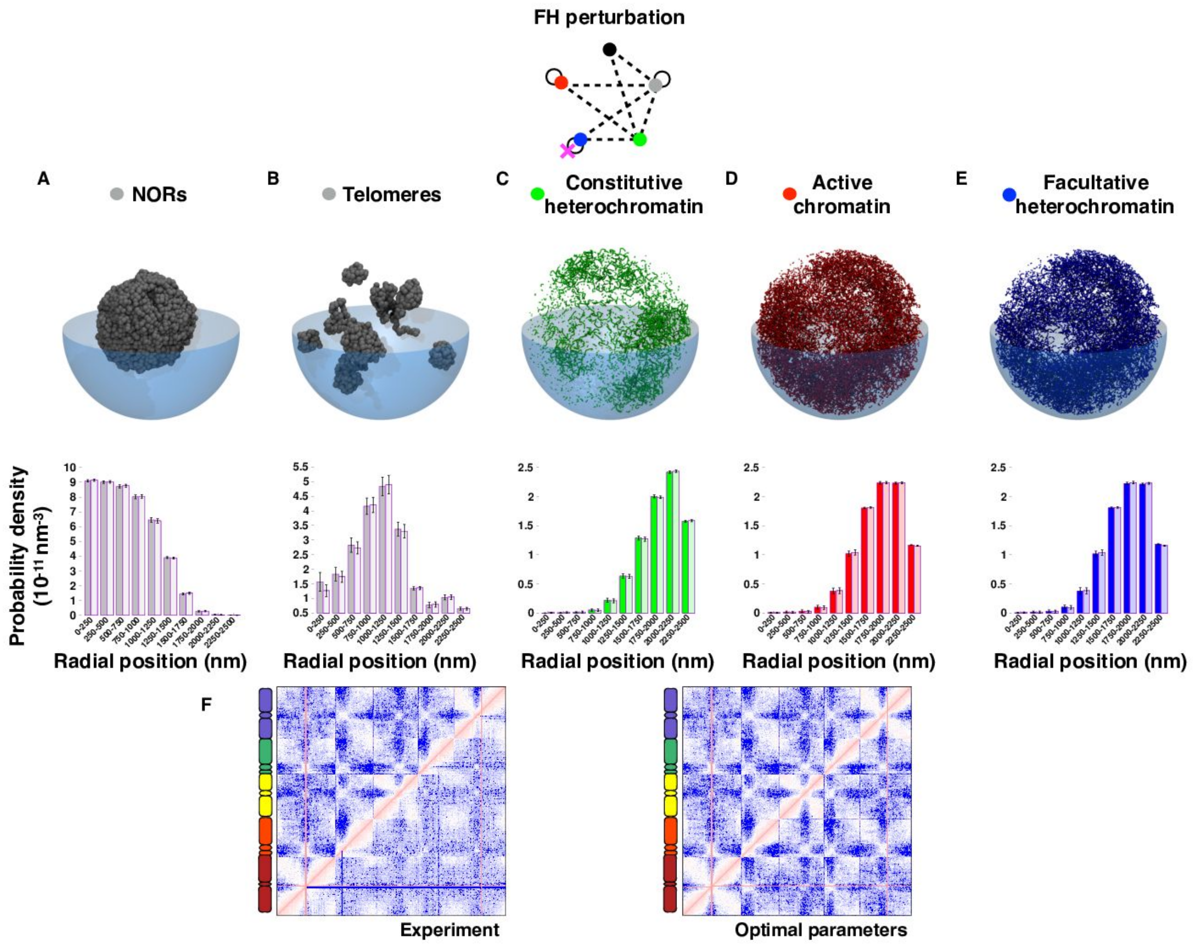


**Supplementary Figure S11. Mild effect of the removal of the self-attraction between facultative heterochromatin (FH) regions.** In the optimal interaction network the FH self-attraction was removed (magenta cross). **(A-E)** (*Top*) Each representative snapshot of the model nucleus shows the 3kbp-regions (beads) of the corresponding chromatin state in the perturbed system at the last frame of one of the 50 simulated trajectory (replicate 1). (*Bottom*) The distribution of the radial positions of the 3kbp-regions in the perturbed systems (dark colour) is compared to the optimal interaction model (light colour) of **Figure 3A**. None significant differences (two-sided Wilcoxon test p-value<0.0001) are detected. **(F)** The genome-wide contact map for the perturbed case (top left triangles) is shown together with (*Left*) the Hi-C and (*Right*) the prediction from the optimal model (bottom right triangles). Removing FH self-attraction does not affect significantly the predicted genome wide maps and the similarity with Hi-C is almost as equivalent as in the optimal model (**Supplementary Figure S7J** and **S8J**).

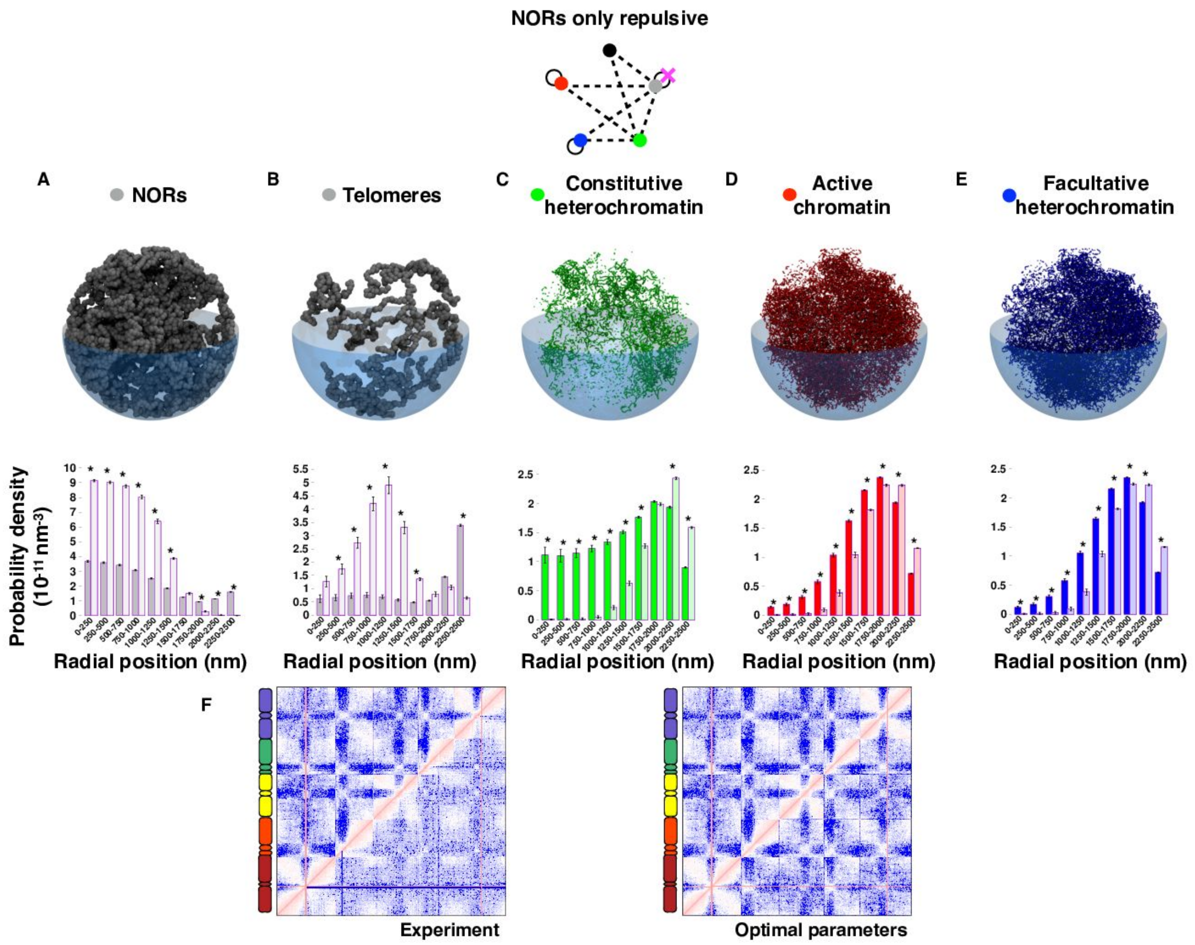


**Supplementary Figure S12. Strong effect of the removal of the self-attraction between nucleolar organising regions (NORs).** In the optimal interaction network the NORs self-attraction was removed (magenta cross). **(A-E)** (*Top*) Each representative snapshot of the model nucleus shows the 3kbp-regions (beads) of the corresponding chromatin state in the perturbed system at the last frame of one of the 50 simulated trajectory (replicate 1). (*Bottom*) The distribution of the radial positions of the 3kbp-regions in the perturbed systems (dark colour) is compared to the optimal interaction model (light colour) of **Figure 3A**. Many significant differences (two-sided Wilcoxon test p-value<0.0001) are detected indicating that a proper compaction of the nucleolus is essential to obtain the expected positioning of each epigenomic state in our models. **(F)** The genome-wide contact map for the perturbed case (top left triangles) is shown together with (*Left*) the Hi-C and (*Right*) the prediction from the optimal model (bottom right triangles). Removing self-attraction between NOR does not affect significantly the predicted genome wide maps and the similarity with Hi-C is almost as equivalent as in the optimal model (**Supplementary Figure S7C** and **S8C**).

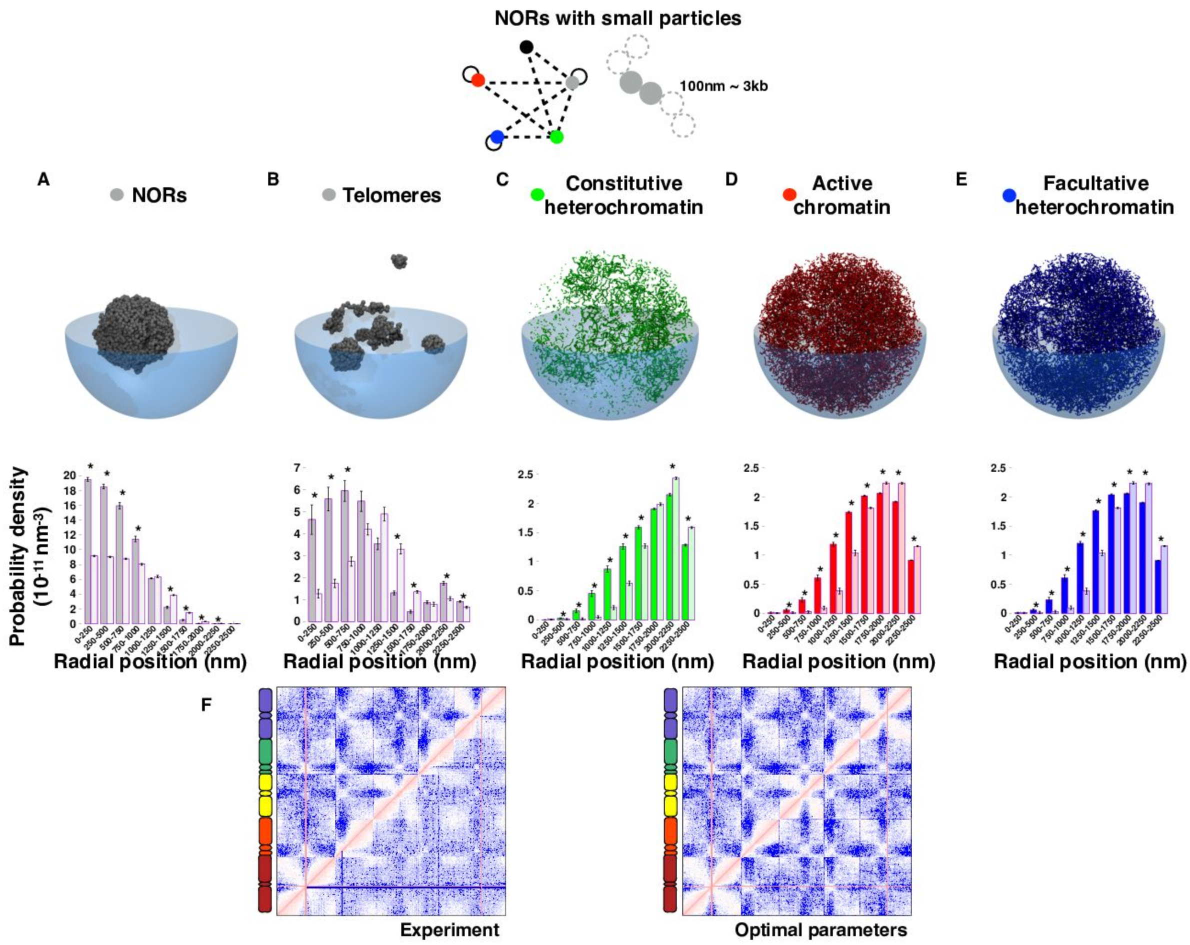


**Supplementary Figure S13. Strong effect of the diameter reduction of nucleolar organising regions (NORs) beads.** Using the optimal interaction network in these simulations the diameter of the NORs beads was decreased from 132 nm to 100 nm (gray cartoon). **(A-E)** (*Top*) Each representative snapshot of the model nucleus shows the 3kbp-regions (beads) of the corresponding chromatin state in the perturbed system at the last frame of one of the 50 simulated trajectory (replicate 1). (*Bottom*) The distribution of the radial positions of the 3kbp-regions in the perturbed systems (dark colour) is compared to the optimal interaction model (light colour) of **Figure 3A**. Many significant differences (two-sided Wilcoxon test p-value<0.0001) are detected indicating that a proper size of the nucleolus is essential to obtain the expected positioning of each epigenomic state in the models. In this perturbed condition, the epigenomic states tend to occupy more central positions. **(F)** The genome-wide contact map for the perturbed case (top left triangles) is shown together with (*Left*) the Hi-C and (*Right*) the prediction from the optimal model (bottom right triangles). Reducing the NOR bead size does not affect significantly the predicted genome wide maps and the similarity with Hi-C is almost as equivalent as in the optimal model (**Supplementary Figure S7D** and **S8D**).

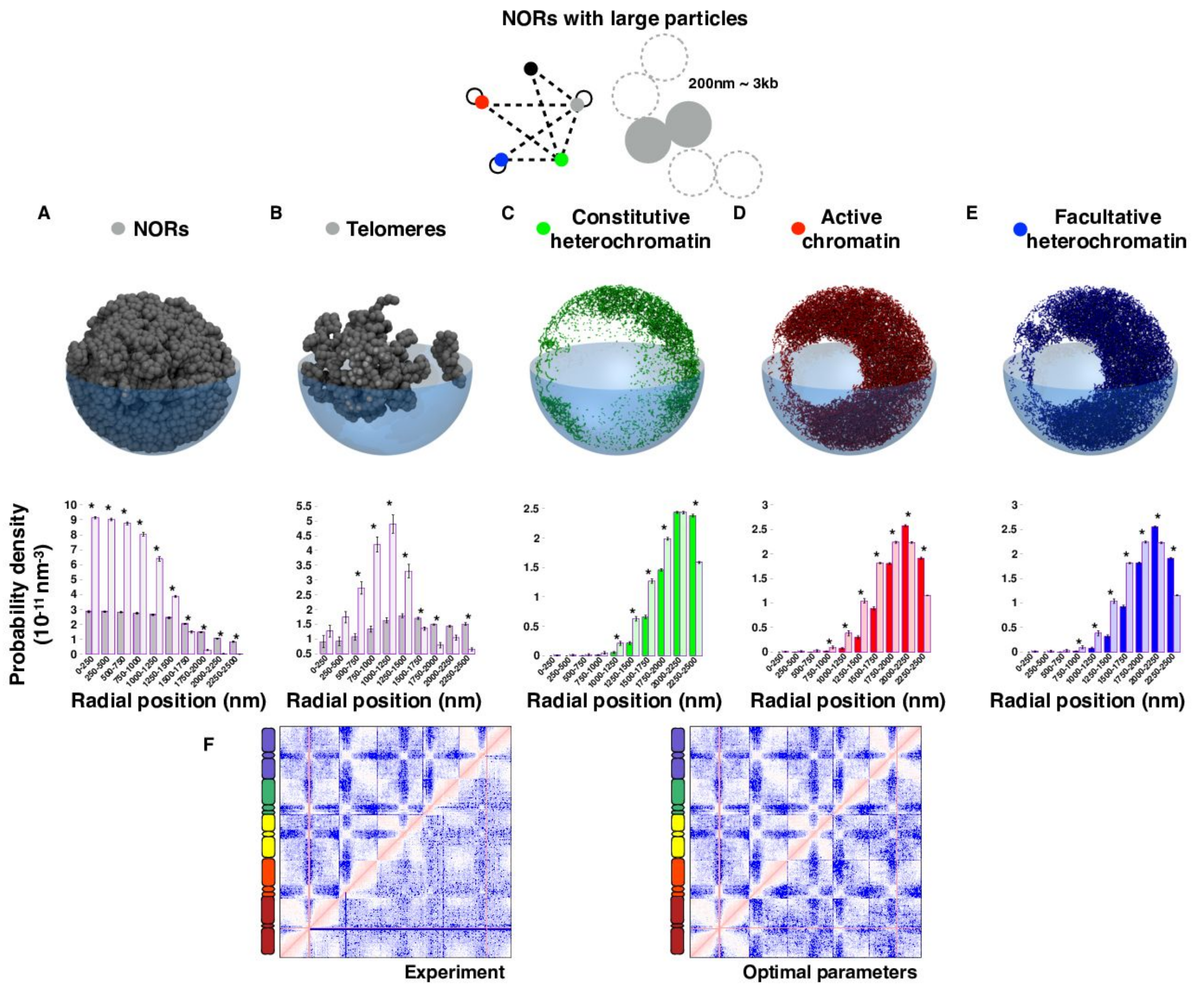


**Supplementary Figure S14. Strong effect of the diameter increase of nucleolar organising regions (NORs) beads.** Using the optimal interaction network in these simulations the diameter of the NORs beads was increased from 132 nm to 200 nm (gray cartoon). **(A-E)** (*Top*) Each representative snapshot of the model nucleus shows the 3kbp-regions (beads) of the corresponding chromatin state in the perturbed system at the last frame of one of the 50 simulated trajectory (replicate 1). (*Bottom*) The distribution of the radial positions of the 3kbp-regions in the perturbed systems (dark colour) is compared to the optimal interaction model (light colour) of **Figure 3A**. Many significant differences (two-sided Wilcoxon test p-value<0.0001) are detected indicating that a proper size of the nucleolus is essential to obtain the expected positioning of each epigenomic state in the models. In this perturbed condition, the epigenomic states tend to occupy more peripheral positions. **(F)** The genome-wide contact map for the perturbed case (top left triangles) is shown together with (*Left*) the Hi-C and (*Right*) the prediction from the optimal model (bottom right triangles). Increasing the NOR bead size does not affect significantly the predicted genome wide maps and the similarity with Hi-C is almost as equivalent as in the optimal model (**Supplementary Figure S7E** and **S8E**).

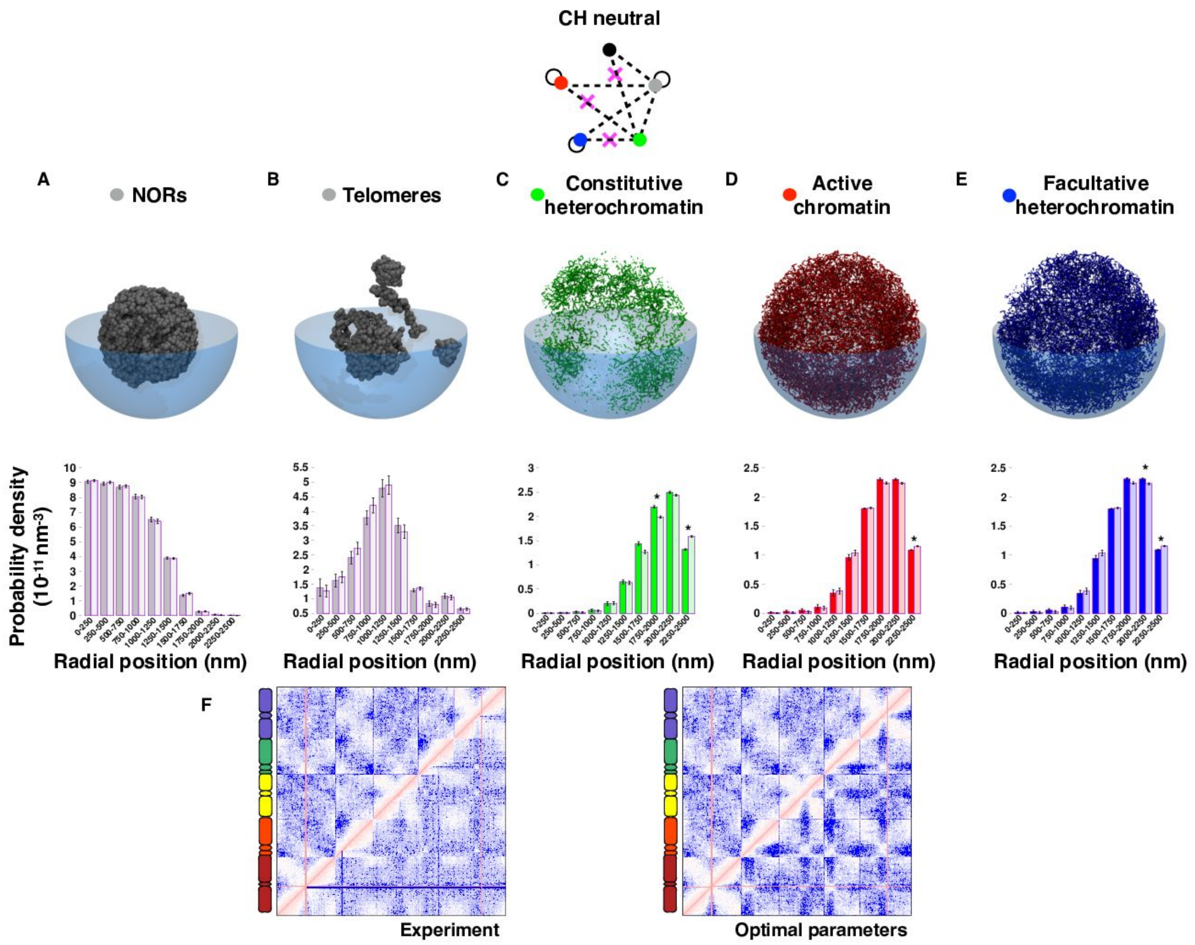


**Supplementary Figure S15. Effect of the removal of the repulsion between constitutive heterochromatin (CH) and other epigenomic states.** In the optimal interaction network the CH repulsion was removed (magenta cross). **(A-E)** (*Top*) Each representative snapshot of the model nucleus shows the 3kbp-regions (beads) of the corresponding chromatin state in the perturbed system at the last frame of one of the 50 simulated trajectory (replicate 1). (*Bottom*) The distribution of the radial positions of the 3kbp-regions in the perturbed systems (dark colour) is compared to the optimal interaction model (light colour) of **Figure 3A**. Few significant differences (two-sided Wilcoxon test p-value<0.0001) were detected indicating that in absence of the repulsive interactions the CH regions lost their preferential peripheral positions. In this perturbed condition, the CH tends to occupy more central positions. **(F)** The genome-wide contact map for the perturbed case (top left triangles) is shown together with (*Left*) the Hi-C and (*Right*) the prediction from the optimal model (bottom right triangles). Removing CH repulsion with other epigenomic states affects significantly the predicted genome wide maps in particular the trans-chromosome areas, although the similarity with Hi-C is almost as equivalent as in the optimal models (**Supplementary Figure S7F** and **S8F**).

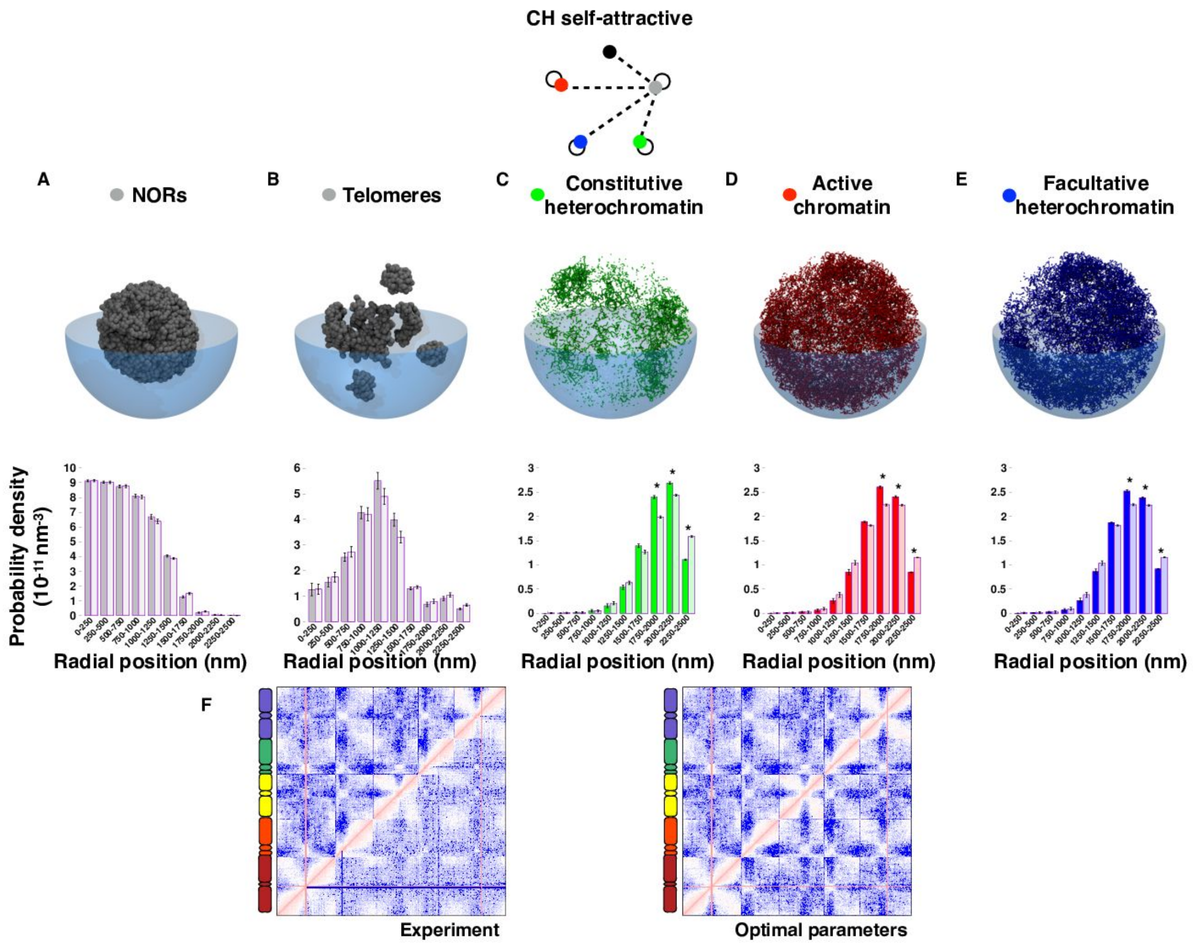


**Supplementary Figure S16. Effect of purely self-attractive constitutive heterochromatin (CH).** In this alternative interaction network, the CH self-attraction (without CH repulsions) was tested. **(A-E)** (*Top*) Each representative snapshot of the model nucleus shows the 3kbp-regions (beads) of the corresponding chromatin state in the perturbed system at the last frame of one of the 50 simulated trajectory (replicate 1). (*Bottom*) The distribution of the radial positions of the 3kbp-regions in the perturbed systems (dark colour) is compared to the optimal interaction model (light colour) of **Figure 3A**. Importantly, few significant differences (two-sided Wilcoxon test p-value<0.0001) were detected indicating that in absence of the repulsive interactions the CH regions lost their preferential peripheral positions. In this perturbed condition, the CH tends to occupy more central positions. **(F)** The genome-wide contact map for the perturbed case (top left triangles) is shown together with (*Left*) the Hi-C and (*Right*) the prediction from the optimal model (bottom right triangles). The purely self-attractive CH scenario does not affect significantly the predicted genome wide maps and the similarity with Hi-C is almost as equivalent as in the optimal model (**Supplementary Figure S7H** and **S8H**).

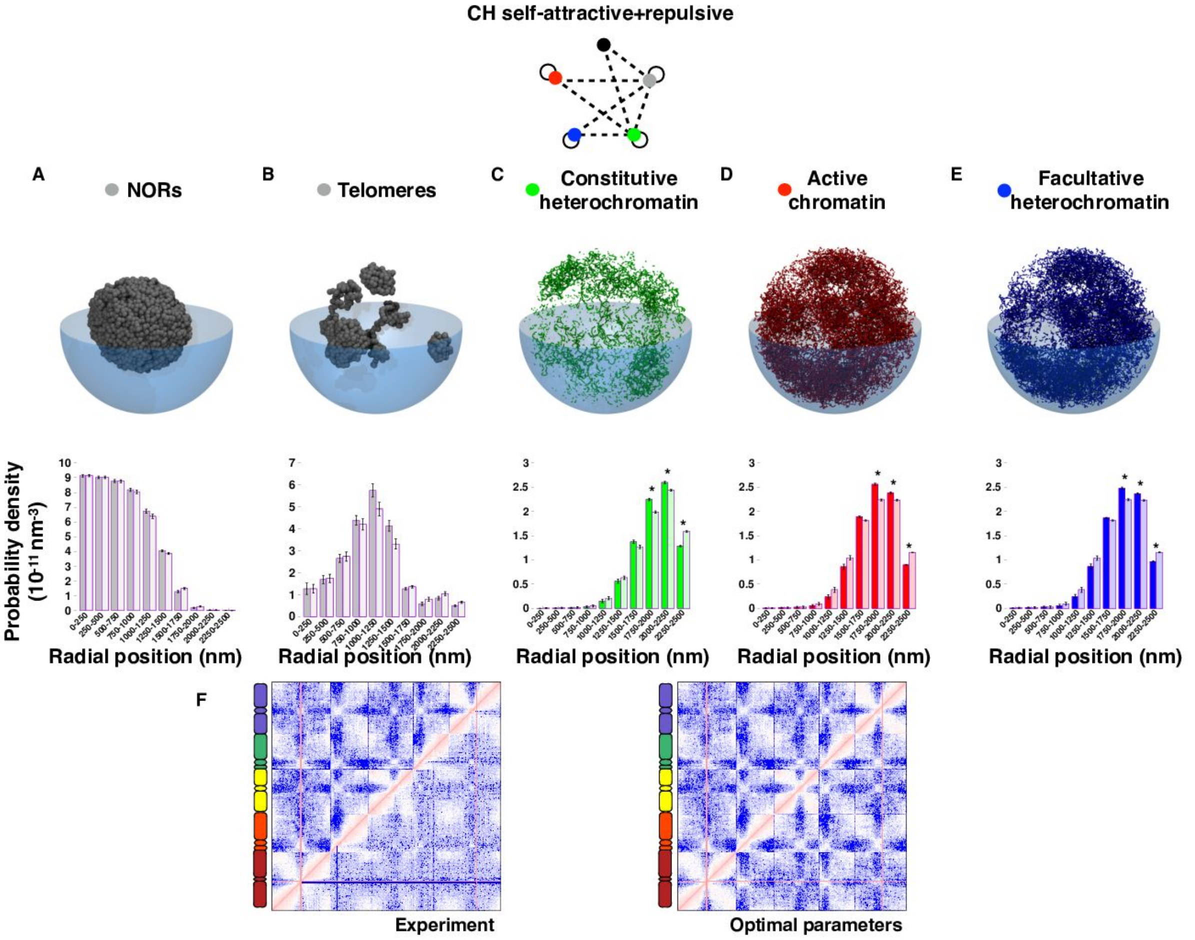


**Supplementary Figure S17. Effect of self-attractive+repulsive constitutive heterochromatin (CH).** In this alternative interaction network, the CH self-attraction combined with CH repulsions was tested. To match the Hi-C compartment strength, the strength of both attractive and repulsive interactions were smaller than the purely attractive and repulsive cases (**Supplementary Figure S4**). **(A-E)** (*Top*) Each representative snapshot of the model nucleus shows the 3kbp-regions (beads) of the corresponding chromatin state in the perturbed system at the last frame of the simulated trajectories. (*Bottom*) The distribution of the radial positions of the 3kbp-regions in the perturbed systems (dark colour) is compared to the optimal interaction model (light colour) of **Figure 3A**. Few significant differences (two-sided Wilcoxon test p-value<0.0001) were detected indicating that the CH repulsive interactions are too mild to favour the CH preferential peripheral positions. In this perturbed condition, the CH, the AC, and the FH states tend to occupy more central positions. **(F)** The genome-wide contact map for the perturbed case (top left triangles) is shown together with (*Left*) the Hi-C and (*Right*) the prediction from the optimal model (bottom right triangles). The self-attractive+repulsive CH scenario does not affect significantly the predicted genome wide maps and the similarity with Hi-C is almost as equivalent as in the optimal model (**Supplementary Figure S7G** and **S8G**).

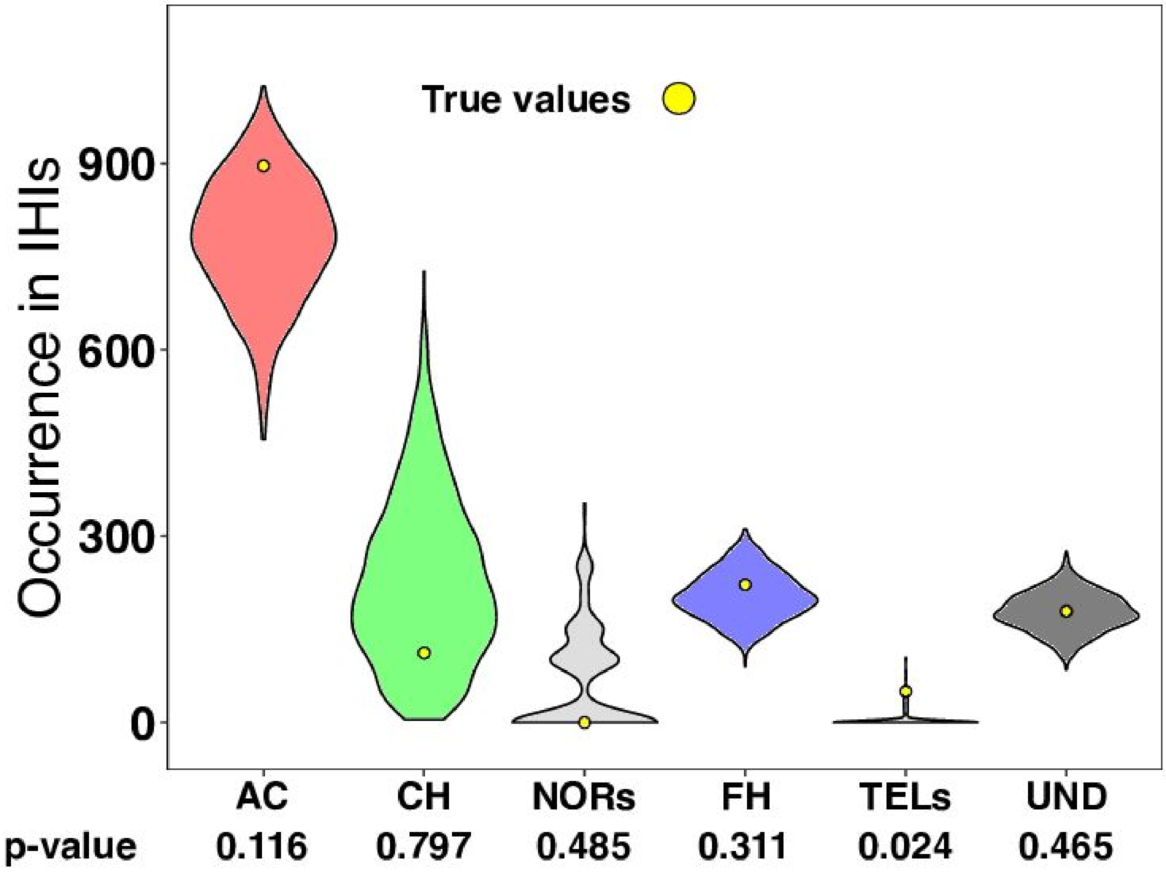


**Supplementary Figure S18**. **Enrichment analysis of (epi)genomic states in KEEs.** The number of beads of each (epi)genomic state (occurrence) in the 10 KNOT engaged elements, KEEs (aka Interacting Heterochromatic Islands (IHIs)) are indicated as yellow points (true values). The violin plots show the distributions of epigenomic-state occurrences in 1,000 randomised KEE-like sets. Briefly, for each of the 10 KEEs of length *l* on chromosome *c*, one region of length *l* was randomly selected on chromosome *c*, and the occurrences of each epigenomic state were computed. Repeating the random sampling N=1,000 times, a distribution of N occurrences per epigenomic state was obtained, and the corresponding distribution was represented as a violin plot. Finally, each p-value quantifies the fraction of times a randomised selection resulted in occurences of the (epi)genomic state higher than the true value. Apart from telomeres, which account for only a few particles, none of the epigenomic states is significantly enriched in KEEs.

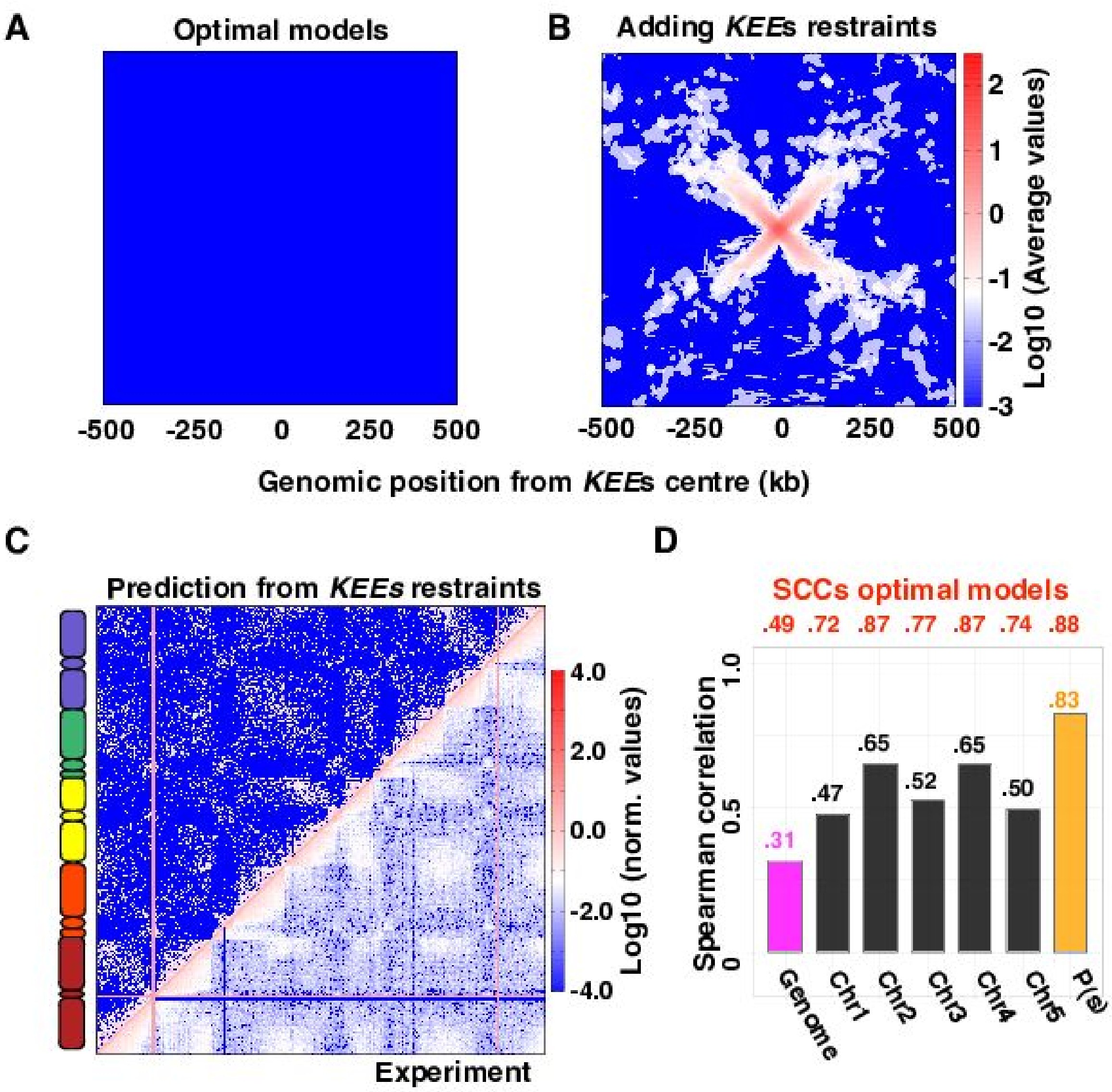


**Supplementary Figure S19**. **Plasticity of the optimal models in response to long-range KEEs-driven restraints. (A-B)** Average contact map of a 1Mbp-region around the KNOT Engaged Elements (KEEs) for the optimal models **(A)** and the KEEs-restrained one **(B)**. **(C)** Genome-wide contact maps in the KEEs-restrained case (top left triangle) and from experiments (bottom right triangle). **(D)** Spearman correlation coefficients (SCCs) obtained comparing the KEEs-restrained models and experiments (Figure 3). The SCC values obtained for the optimal interaction model are given in red.

### SUPPLEMENTARY METHODS

**Epigenomic states analyses.** From [2,4], the (ChIP-seq) epigenomic data of 4 chromatin marks were collected at a 400bp-resolution: H3K4me2, H3K4me3 signatures of active genes, H3K27me3 signature of facultative, polycomb-like heterochromatin, and H3K9me2 specific to constitutive heterochromatin. To each 3kbp genomic region, an average epigenomic signal was assigned for each of these 4 marks. A K-means algorithm (Euclidean distance, k=4) allowed clustering the 3kbp-regions into 4 groups: active chromatin (AC) enriched in H3K4me2/3, facultative heterochromatin (FH) enriched in H3K27me3, constitutive heterochromatin (CH) enriched in H3K9me2 and undetermined (UND) depleted in all of these 4 marks. The lengths and the genomic localization of the domains assigned to each of the 4 chromatin states were used to generate the plots in **Figure 1B-C** and in **Supplementary Figure S1**.

**The chromosome polymer model.** Molecular dynamics simulations of the diploid *Arabidopsis thaliana* genome were run using the 30nm-fibre model [6], in which each bead of unitary mass (m=1.0) hosts 3 kilobase pairs (kbp) of DNA sequence and has a diameter of σ=30 nm. Each *A. thaliana* chromosome was represented as a chain of beads using the Kremer-Grest bead-spring model [7] with the same parameter as in Ref. [8]:

$$H = U_{EV} + U_C + U_{BEND} \quad (1)$$

The first term was a truncated and shifted Lennard-Jones potential that controls the *cis*- and *trans*-chromosome excluded volume interactions:

$$U_{EV}(i,j) = \begin{cases} 4k_BT\epsilon_{ij}\left[\left(\frac{\sigma}{d_{i,j}}\right)^{12} - \left(\frac{\sigma}{d_{i,j}}\right)^{6} + 1/4\right] & \text{if } d_{i,j} \leq 2^{1/6}\sigma, \\ 0 & \text{if } d_{i,j} > 2^{1/6}\sigma. \end{cases} \quad (2)$$

where $k_B$ is the Boltzmann constant, T the temperature, $\varepsilon_{i,j}$ is equal to 10 if $|i-j| = 1$, and 1 otherwise, $\sigma = 30\ nm$ was the thickness of the chain and $d_{i,j}$ is the modulus of $\overline{d_{i,j}} = \overline{r_i} - \overline{r_j}$, that is the distance vector between the monomers *i* and *j* at positions $\overline{r_i}$ and $\overline{r_j}$, respectively. The second term was a FENE potential that ensures the chain connectivity between consecutive beads on the same polymer chain:

$$U_C(i, i+1) = -0.5KR_0^2 ln\left[1 - \left(\frac{d_{i,i+1}}{R_0}\right)^2\right]$$

where K=0.33 $k_BT/nm^2$ and $R_0$=45 nm. The combined action of the connectivity and excluded volume interaction between consecutive beads was such that the average bond length was close to 30 nm and never exceeded 40 nm.

The third term is a (Kratky-Porod) bending potential:

$$U_{BEND} = \frac{k_B T l_p}{\sigma}\left(1 - \frac{\overline{d_{i,i+1}}\cdot\overline{d_{i+1,i+2j}}}{d_{i,i+1}d_{i+1,i+2j}}\right)$$

where the chain persistence length, $l_p$, has been set equal to 150 nm to reproduce the experimental bending properties of the chromatin fibre [9]. The lengths of *A. thaliana* chromosomes (in base-pairs and in models' beads) and the genomic locations of special sequences (NORs and centromeres) are reported in **Supplementary Table S1**.

The dynamics of the polymer model was simulated using the LAMMPS simulation package (version 31Mar2017) [10] integrating the (underdamped) Langevin equation of motion:

$$m\ddot{r}_{i\alpha} = -\partial_{i\alpha}H - \gamma\dot{r}_{i\alpha} + \eta_{i\alpha}(t)$$

where m is the mass of the bead that was set equal to the LAMMPS default value, H is the energy of the system in Eq. 1, the index *i* runs over all the particles in the system, and $\alpha = (x, y, z)$ indicates the Cartesian components, and $\gamma = 0.5\ \tau_{LJ}^{-1}$ is the friction coefficient with $\tau_{LJ} = \sigma(m/\varepsilon)^{1/2}$ is the Lennard-Jones time. The stochastic term $\eta_{i\alpha}$ satisfies the fluctuation-dissipation conditions. The integration time step used in the numerical integration was equal to $\Delta t = \alpha\ \tau_{LJ}$, where the factor α was adapted, as specified below, to the different stages of the preparation and production runs

**Epigenomics-driven interactions**. In this study, additional epigenomics-based short-range interactions were used to test how the attractions or repulsions between the *A. thaliana* chromatin states shape the chromosome organization. These interactions have been modelled using attractive or repulsive Lennard-Jones potentials of various strengths (see Eq. 2). with the following parameters:

- Repulsive interactions: σ=90 nm, cutoff= $\sqrt[6]{2}\sigma$, and strengths ε in $k_BT$ as reported in **Supplementary Figures S1** and **S2**. This acted as a purely repulsive Lennard-Jones where the particles fell each other as thicker (45 nm radius) than their nominal radius (15 nm).
- Attractive interactions: σ=30 nm, cutoff= $2.5\sigma$ =75 nm, and strengths ε in $k_BT$ as reported in **Supplementary FIgure S2** and **S3**. The cutoff allowed to include the attractive part of the Lennard-Jones potential.

**Single chromosome simulations.**

To identify the optimal epigenomics-based interaction, the simulations were first applied to study the folding of chromosome 4 (Chr4), which is the shortest chromosome of the *A. thaliana* karyotype. Chr4 was described as a chain of 6,195 beads (excluding the NOR4 to simplify) which could be properly taken into account only in genome-wide simulations (**Supplementary Table S1**). To enhance the statistics per run, a toy system was produced where 5 copies of Chr4 were placed in a cubic simulation box of side equal to 2.76 μm with periodic boundary conditions. The overall system density was therefore about 0.004 bp/nm$^3$ as that corresponds to the DNA density of the *A. thaliana* genome [8,11]. To mimic a mitotic-like state, each model chromosome was initially prepared in an elongated solenoidal-like configuration [11], and the 5 copies were placed in a random, yet non-overlapping arrangement inside the cubic simulation box. Dynamics of 10 independent trajectories were then simulated. To remove any excessive intra-chain strain of the orderly designed mitotic arrangement and to relax the chromosome conformations from the schematic rosette re-arrangement, the chromosome models were evolved for 10,000$\tau_{LJ}$ (10,000,000 $\Delta$t with $\Delta$t=0.001 $\tau_{LJ}$), to obtain partially de-condensed arrangements.

After applying the epigenomic-driven interactions, the chromosomes were energy-minimized using the Polak-Ribiere version of the conjugate gradient algorithm [10] (Lammps command: minimize 1.0e-4 1.0e-6 100000 100000) and simulated for 10 $\tau_{LJ}$ (10,000 $\Delta$t with $\Delta$t=0.001 $\tau_{LJ}$) to favour the mild adaptation of the polymers to the newly introduced interactions.

Using the steps above, 10 independent initial conformations were obtained. Each of them was then simulated for 120,000 $\tau_{LJ}$ (20,000,000 $\Delta$t with $\Delta$t=0.006 $\tau_{LJ}$), which corresponds to a few hours [11]. From these simulations, starting from 6,000 $\tau_{LJ}$ and collecting one conformation every 3,000 $\tau_{LJ}$ of dynamics, a total of 39 snapshots per replicate were stored. The resulting set of 390 out-of-equilibrium conformations was used for the downstream analysis.

**Preparation of the genome-wide initial conformations.** To test how the initial mitotic-like chromosome arrangements affect the accuracy of the obtained models, three different chromosome shapes were generated: (i) linear rod-like chromosomes, (ii) V-shaped chromosomes generated by linear pulling along the same direction, (iii) V-shaped chromosomes generated by linear pulling along radial directions.

**Generation of the linear chromosome arrangements**. The generation of the initial linear chromosome arrangements is illustrated in **Supplementary Video S2**. The detailed account of the steps is presented in the following paragraphs. As in the single-chromosome system, the chromosomes were initially generated as cylindrical rods made of stacked rosettes [11,12]. The initial positions and the orientations of the rods were chosen inside a sphere of radius 2.5 μm centred in the origin O=(0,0,0) of the system. This confinement was enforced as a rigid wall with a 12/6 Lennard-Jones potential and mimicked the typical shape and size of *A. thaliana* nuclei [13]. Such positioning was done starting from the longest chromosome pair down to the shortest one (**Supplementary Table S1**) and was biased such that the regions hosting the nucleolar organizing regions (NORs, see **Supplementary Table S1**) were drawn inside a sphere of radius 450 nm and centred in O. The remaining chromosome pairs (1, 3, and 5) were drawn with random positions and orientations inside the nuclear sphere, with the condition that none of their particles could occupy a sphere centred in O and of radius 600 nm. This particular strategy was designed to favour the formation of the nucleolus in the following preparation stages. After an energy minimization (minimize 1.0e-4 1.0e-6 100000 100000), the rod-like chromosome shapes were pinned by 35,052 harmonic bonds maintaining loops of ~42 kbp (14 beads) within planar rosettes and between contiguous rosettes. The equilibrium distance of each harmonic bonds was set to $d_{eq}$=60 nm and the spring constant was ramped up from 0 to 1.1 $k_BT/nm^2$ during a run of 10 $\tau_{LJ}$ (10,000 $\Delta t$ with $\Delta t$ = 0.001 $\tau_{LJ}$) (representative snapshot in **Supplementary Figure S6A**). Next, during a run of 1,200 $\tau_{LJ}$ (200,000 $\Delta t$ with $\Delta t$ = 0.006 $\tau_{LJ}$), the harmonics pinning the looped structure were maintained stable (K=1.1 $k_BT/nm^2$ and $d_{eq}$=60 nm) for all beads but NORs and telomeres, and a Lennard-Jones self-attractive interaction ($\sigma$=30 nm, $\varepsilon$=1.0 $k_BT$, and cutoff=75 nm) between NORs and telomeres beads was switched on to favour the formation of a single nucleolus. After releasing all the harmonic bonds, a further run of 6,000 $\tau_{LJ}$ (1,000,000 $\Delta t$ with $\Delta t$ = 0.006$\tau_{LJ}$) was applied to finalise the nuclear formation. Once the NORs particles were grouped to form a single globule, the system was evolved to inflate the NORs and telomeres particles to reach a radius of 66 nm such that the expected total volume of these particles accounted for the volume of a sphere of radius 1.4 µm (the typical radius of observed nucleolus in *A. thaliana*) under the hypothesis of Random close packing (64% of the total volume of the sphere is actually occupied by beads).

$$0.64 V_{Nucleolus} = V_{Beads}$$

$$0.64 \tfrac{4}{3}\pi R^3_{Nucleolus} = (N_{NORs} + N_{TELs}) \tfrac{4}{3}\pi R^3_{Bead}$$

$$R_{Bead} = \sqrt[3]{\frac{0.64}{N_{NORs}+N_{TELs}}} R_{Nucleolus} = \sqrt[3]{\frac{0.64}{5062+1000}} 1400\ nm \simeq 66\ nm$$

The inflation of the NORs and telomeres particles was done by 17 short simulations of 0.1 $\tau_{LJ}$ (1,000 $\Delta t$ with $\Delta t$=0.0001 $\tau_{LJ}$) each, in which the radius of each NORs and telomeres bead was increased from 15 nm to 66 nm in steps of 3 nm adapting connectivity and excluded volume interactions. The system was then simulated for 3,000 $\tau_{LJ}$ (1,000,000 $\Delta t$ with $\Delta t$ = 0.003 $\tau_{LJ}$) to stabilize the formation of the nucleolus. Before applying the specific epigenomics-driven interactions the system was further relaxed for 10,000 $\tau_{LJ}$ (10,000,000 $\Delta t$ with $\Delta t$ = 0.001 $\tau_{LJ}$).

**Generation of the v-shaped chromosome arrangements**. To generate v-shaped chromosomes, the chromosomes arranged as the one in **Supplementary Figure S6A** were pulled simultaneously to bend their conformations by the respective centromere, that in *A. thaliana* coincides with the kinetochore [14,15]. Two pulling protocols were applied.

**V-shape from parallel pulling**. The chromosomes were pulled along the same (parallel) direction y=(0.0,1.0,0.0) (**Supplementary Figure S6B**) as illustrated in the **Supplementary Video S3**. The pulling was done from the conformations exemplified in **Supplementary Figure S6A** by translating rigidly all chromosomes 4.5 µm along the -y=(0.0,-1.0,0.0) direction and, then, by tethering the centre of mass of each centromere to its initial (untranslated) positions with a harmonic spring (K=0.05 $k_BT/nm^2$ and $d_{eq}$=0 nm) imposed with a run of 200 $\tau_{LJ}$ (200,000 $\Delta$t with $\Delta t=0.001\tau_{LJ}$). During the pulling runs, the harmonic bonds (K=1.1 $k_BT/nm^2$ and $d_{eq}$=60 nm) were maintained. Next, the harmonic bonds pinning the stacked rosettes were kept in place (K=1.1 $k_BT/nm^2$ and $d_{eq}$=60 nm), while the beads of the NORs and the telomeres (see **Supplementary Table S1**) were left free to de-condense from their rod-like arrangement and to condense between each other to form the nucleolus. The formation of the nucleolus was implemented applying a Lennard-Jones attraction ($\sigma$=30 nm, $\varepsilon$=1.0 $k_BT$, and cutoff=75 nm) in two separated runs of 600 $\tau_{LJ}$ (100,000$\Delta$t with $\Delta t$ = 0.006 $\tau_{LJ}$). Firstly, it was applied only between NORs, and secondly, it was extended also within NORs. These two runs were designed such that the NORs beads could explore the space around them, interact with the other close NORs regions, and finally condense altogether. To enforce spherical nuclear confinement, the first 60 $\tau_{LJ}$ of these runs were used to compress the system inside the nuclear sphere changing the spherical radius from 4.0 µm to the target 2.5 µm [13,16,17]. Next, while all the harmonic bonds were released, the system was relaxed for 6,000 $\tau_{LJ}$ (1,000,000 $\Delta$t with $\Delta t$ = 0.006 $\tau_{LJ}$). During this run, the NORs particles continued their condensation and group in a globule in the central volume of the nucleus, while the other beads de-condensed to occupy a large portion of the nuclear volume. Once the NORs particles were grouped to form a single globule, the system was evolved to inflate the NORs and telomeres particles to reach a radius of 66 nm as in the linear case. The inflation of the NORs and telomeres particles was done by 17 short simulations of 0.1 $\tau_{LJ}$ (1,000 $\Delta$t with $\Delta t$ = 0.0001 $\tau_{LJ}$) each, in which the radius of the NORs and TEL particles was increased from 15 nm to 66 nm in steps of 3 nm. Similar strategies were used to obtain NORs particles of diameters 100 nm and 200 nm in the variant simulations. Before applying the epigenomic-driven interactions, the system was relaxed with two runs: the first of 3,000 $\tau_{LJ}$ (1,000,000 $\Delta$t with $\Delta t$ = 0.003 $\tau_{LJ}$). and the second of 10,000 $\tau_{LJ}$ (10,000,000 $\Delta$t with $\Delta t$ = 0.001 $\tau_{LJ}$).

**V-shape from radial pulling**. Starting from the linear conformations (see **Supplementary Figure S6A**), each chromosome was pulled along different radial directions as illustrated in the **Supplementary Video S4**. The pulling was done from the conformations exemplified in **Supplementary Figure S6A** by applying to each centromere a harmonic spring (K=0.05 $k_BT/nm^2$ and $d_{eq}$=5.5 μm) to push away the centromeres along the direction joining the origin of the system and the centre of mass of each centromere (**Supplementary Figure S6C**) with a run of 100 $\tau_{LJ}$ (100,000 $\Delta t$ with $\Delta t$=0.001 $\tau_{LJ}$). During these runs, the harmonic bonds (K=1.1 $k_BT/nm^2$ and $d_{eq}$=60 nm) were maintained. Next, the centromeres were evolved as rigid bodies for 100 $\tau_{LJ}$ (100,000$\Delta t$ with $\Delta t$=0.001$\tau_{LJ}$) during which the nuclear confinement was enforced with a sphere of changing radius from 4.0 μm to the target 2.5 μm [13,16,17]. Next, the system was relaxed for 6,000 $\tau_{LJ}$ (1,000,000 $\Delta t$ with $\Delta t$ = 0.006 $\tau_{LJ}$) by releasing all the harmonic bonds. During this run, the NORs particles condensed in a globule in the central volume of the nucleus, while the other beads de-condensed to occupy a large portion of the nuclear volume. Once the NORs particles were grouped to form a single globule, the system was then evolved to inflate the NORs and telomeres particles to reach a radius of 66 nm as in the linear case. The inflation of the NORs and telomeres particles was done by 17 short simulations of 0.1 $\tau_{LJ}$ (1,000 $\Delta t$ with $\Delta t$ = 0.0001 $\tau_{LJ}$) each in which the radius of the NORs and TEL particles was increased from 15 nm to 66 nm in steps of 3 nm. Finally, the system was relaxed with two runs: the first of 6,000 $\tau_{LJ}$ (2,000,000 $\Delta t$ with $\Delta t$ = 0.003 $\tau_{LJ}$) and the second of 6,000 $\tau_{LJ}$ (1,000,000 $\Delta t$ with $\Delta t$ = 0.006 $\tau_{LJ}$).

**Genome-wide chromosome simulations.** Using the strategy presented above, 50 replicates systems were obtained for each of the 3 chromosome shapes using LAMMPS. On these systems, simulations using the epigenomics-based interactions were performed. Short energy-minimization (minimize 1.0e-4 1.0e-6 100000 100000) and simulation run 100 $\tau_{LJ}$ (100,000 $\Delta t$ with $\Delta t$ = 0.001 $\tau_{LJ}$) preceded the production runs that lasted for 120,000 $\tau_{LJ}$ (20,000,000 $\Delta t$ with $\Delta t$ = 0.006 $\tau_{LJ}$). During this long simulation, starting from 6,000 $\tau_{LJ}$, (1,000,000 $\Delta t$ with $\Delta t$ = 0.006 $\tau_{LJ}$), one conformation every 3,000 $\tau_{LJ}$ (500,000 $\Delta t$ with $\Delta t$ = 0.006 $\tau_{LJ}$) of dynamics was stored for a total of 39 snapshots per replicate. The resulting 1,950 out-of-equilibrium conformations were used for the downstream analysis.

**KEEs-driven steered molecular dynamics**. For the analysis presented in **Figure 7**, 50 steered molecular dynamics simulations were performed starting from the final snapshots of the optimal-model simulations to promote the spatial proximity between the closest KNOT ENGAGED ELEMENTS (KEEs), aka IHIs [18,19]. Specifically, the distances between the 10 pairs of KEEs (central beads) were computed both cis- and trans-chromosome in all the 50 initial conformations and the closest bead pairs across all the snapshots were selected to be co-localized using steered molecular dynamics. In particular, the percentages of co-localized KEEs pairs in the models were equal to the experimental FISH association rates for the 4 KEEs pairs experimentally probed in Ref. [18] (20% of the KEE6-KEE1, 35% of the KEE5-KEE4, 66% of the KEE6-KEE3, and 16% of the KEE5-KEE10 pairs), and to the average rate of 34% for all the other KEEs pairs. The KEEs pairs contacts were enforced with harmonic constraints (K ramping from 0 to 0.01 $K_BT/nm^2$ and $d_{eq}$=0 nm) between the KEEs during 50 runs (one per replicate) of 12,000 $\tau_{LJ}$ (2,000,000 $\Delta t$ with $\Delta t$ = 0.006 $\tau_{LJ}$) each.
